# Ancient genomic linkage couples metabolism with erythroid development

**DOI:** 10.1101/2023.09.25.558944

**Authors:** Alexandra E. Preston, Joe N. Frost, Mohsin Badat, Megan Teh, Andrew E. Armitage, Ruggiero Norfo, Sarah K. Wideman, Muhammad Hanifi, Natasha White, Noémi Roy, Bart Ghesquiere, Christian Babbs, Mira Kassouf, James Davies, Jim R. Hughes, Rob Beagrie, Douglas R. Higgs, Hal Drakesmith

## Abstract

Generation of mature cells from progenitors requires tight coupling of differentiation and metabolism. During erythropoiesis, erythroblasts are required to massively upregulate globin synthesis then clear extraneous material and enucleate to produce erythrocytes^1–3^. *Nprl3* has remained in synteny with the α-globin genes for >500 million years^4^, and harbours the majority of the α-globin enhancers^5^. Nprl3 is a highly conserved inhibitor of mTORC1, which controls cellular metabolism. However, whether Nprl3 itself serves an erythroid role is unknown. Here, we show that Nprl3 is a key regulator of erythroid metabolism. Using Nprl3-deficient fetal liver and adult competitive bone marrow - fetal liver chimeras, we show that NprI3 is required for sufficient erythropoiesis. Loss of Nprl3 elevates mTORC1 signalling, suppresses autophagy and disrupts erythroblast glycolysis and redox control. Human CD34+ progenitors lacking NPRL3 produce fewer enucleated cells and demonstrate dysregulated mTORC1 signalling in response to nutrient availability and erythropoietin. Finally, we show that the α-globin enhancers upregulate *NprI3* expression, and that this activity is necessary for optimal erythropoiesis. Therefore, the anciently conserved linkage of *NprI3*, α-globin and their associated enhancers has enabled coupling of metabolic and developmental control in erythroid cells. This may enable erythropoiesis to adapt to fluctuating nutritional and environmental conditions.

## Main

Erythropoiesis produces ∼2 million erythrocytes per second in humans^2^, representing the highest synthesis of all cell types^6^, likely the largest continuous metabolic challenge of the body. Erythropoiesis also involves tightly controlled transcriptional regulation of the α- and β-globin genes, ensuring that globin synthesis occurs in a timely and sufficient manner. The regulation of α-globin expression by its enhancers has been extensively characterised^7–11^. Interestingly, α-globin has been maintained in synteny with Nitrogen permease regulator-like 3 (*Nprl3*) since before the divergence of jawed (gnathostome) and jawless (agnathan) vertebrates, and before the teleost-specific genome duplication. The α- and β-globin gene clusters separated in amniote vertebrates due to transposition of the proto β-globin gene^12^. However, *Nprl3*, α-globin and their enhancers have been conserved in synteny for >500 million years, leading us to hypothesise that their co-localisation underlies a functionally significant partnership between these genomic elements.

*Nprl3* is conserved throughout Animalia, Fungi, Excavata, SAR (Stramenopiles, Alveolates, Rhizaria) and Amoebazoa; the only taxon that evidently lacks an *Nprl3*-like gene is that of green algae and land plants^12^. *Nprl3* is broadly expressed across vertebrate cell types, and its loss is embryonic lethal^13^. It is a member of the GATOR-1 complex, which negatively regulates mTORC1. mTORC1 is a central metabolic controller, promoting anabolic processes (such as protein translation and synthesis^14^), inhibiting catabolism (such as autophagy^15^), and influencing metabolic pathways of ATP production^16^. Generally, anabolic metabolic outputs require active mTORC1, whereas catabolic signals occur under low mTORC1 signalling. mTORC1 serves critical roles in erythroid cells, contributing to an essential anabolic - catabolic balance during erythropoiesis: erythroblasts undergo rapid haemoglobinisation followed by high autophagic and proteasomal activity to expel or degrade organelles, ultimately resulting in a cytosol comprising 98% haemoglobin^17,18^. Nevertheless, how mTORC1 is dynamically regulated during this process remains poorly understood, and to the best of our knowledge, the role of Nprl3 in erythropoiesis has not been explored.

*Nprl3* introns contain 4 of the 5 α-globin enhancers in mice, and 3 of 4 in humans (Fig. 1a). Furthermore, published data show that *Nprl3* RNA expression increases during erythroid commitment^19^. Considering its conserved genomic context and function regulating mTORC1, we hypothesised that Nprl3 would serve a key role in erythroid metabolism, potentially transcriptionally supported by its location in a transcriptional hub, thus linking metabolism with completion of the erythroid programme during red blood cell synthesis.

**Figure 1.**
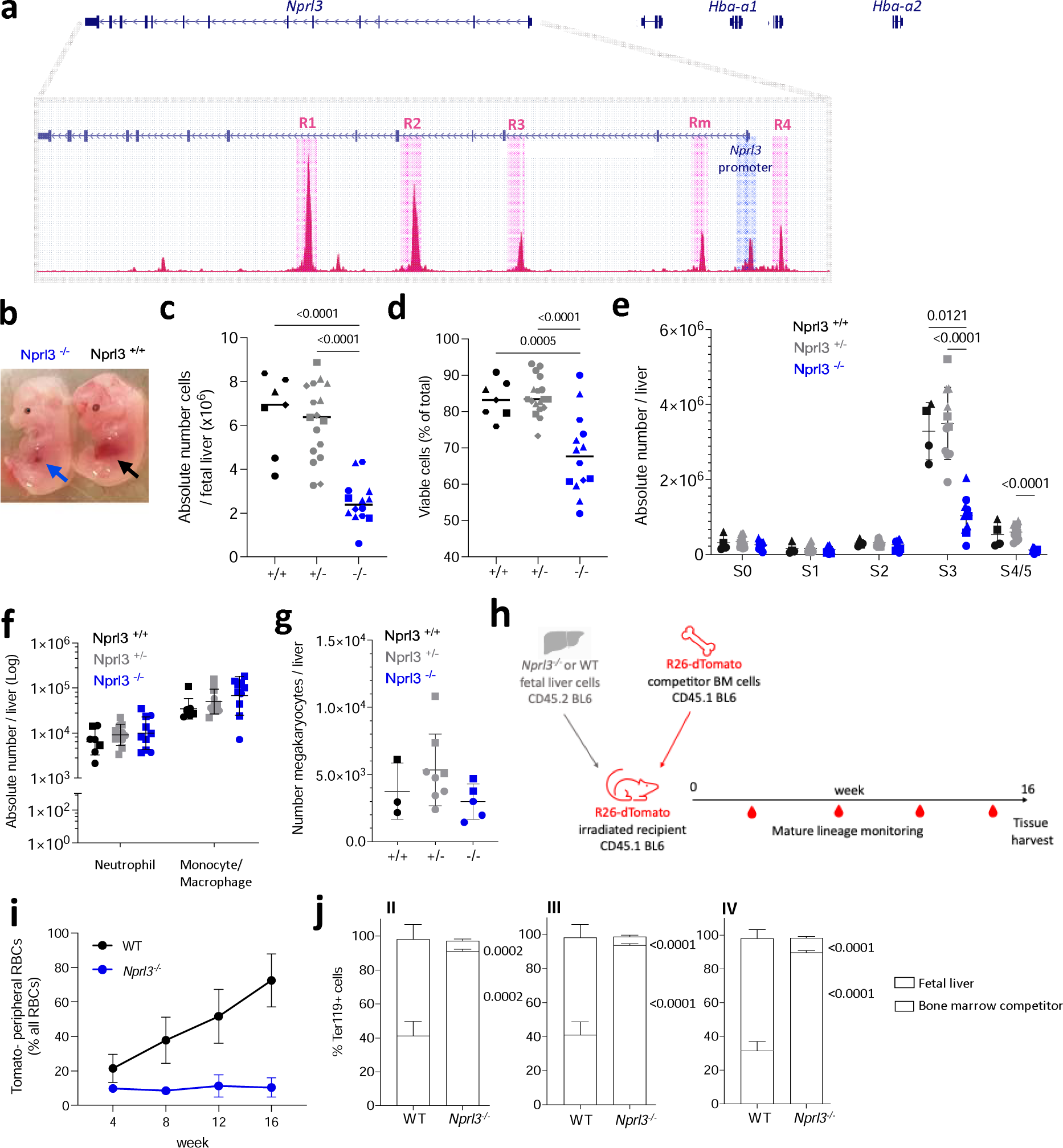
Loss of Nprl3 impairs erythroid differentiation in murine fetal liver and adult competitive chimaeras. a) Schematic illustrating the relative genomic localisation of Nprl3 and Hba-1/2, as well as the positions of the 5 murine α-globin enhancers, MCS-R1, R2, R3, Rm and R4, defined here by ATAC-Seq (GSE174110). b) Representative images of E13.5 Nprl3^-/-^ and Nprl3^+/+^ embryos. c) Total number and d) viability of cells per fetal liver according to genotype (n = 5 litters). Data are expressed as the mean ± SD. Analysed by One-way ANOVA followed by Tukey’s test. e) Absolute number of erythroid cells in stages S0-S5 of differentiation in fetal livers of Nprl3^+/+^, Nprl3^+/-^ and Nprl3^-/-^ embryos (n = 3 litters). S0/S1, CFU-Es; S2, proerythroblasts; S3, basophilic erythroblasts; S4/S5, polychromatic erythroblasts - enucleated cells. Points indicate individual embryos. Data are expressed as the mean ± SD. Analysed by Two-way ANOVA followed by Tukey’s test. f) Absolute numbers of CD11b+ Ly6G+ neutrophils and CD11b+ Ly6G-F4/80+ monocytes/macrophages (n = 3 litters). Data have been log-transformed for visualization and statistical analysis. Each point represents an individual embryo, and the geometric mean ± geometric SD are indicated. Analysed by Two-way ANOVA followed by Tukey’s test. g) Absolute number of c-kit-CD41+ ‘megakaryocyte’ cells per fetal liver (n = 2 litters). Data are expressed as mean ± SD, and analysed by One-way ANOVA followed by Tukey’s test. h) Schematic illustration of the competitive chimaera experimental design. i) Proportion of peripheral RBCs derived from WT and Nprl3^-/-^ fetal liver cells. j) Competitivity comparison showing the proportions of cells in each erythroblast stage (defined by flow cytometry using CD44 and forward scatter: II, basophilic; III, polychromatic, and IV, orthochromatic erythroblasts) derived from fetal liver vs. competitor bone cells, according to the genotype of the transplanted fetal liver. Analysed in recipient bone marrow at week 16 post-transplantation. Data analysed by One-way ANOVA followed by k’s test (n = 8 mice per group). Upper p-value compares fetal liver, lower compares bone marrow competitors. Bars plotted at the mean, error bars indicate standard deviation.

## Nprl3-deficient erythropoiesis is ineffective in fetal and adult mice

We employed an *Nprl3* constitutive knockout model (*Nprl3*^-/-^), in which the entire *Nprl3* promoter region is deleted, leaving the α-globin enhancers (MCS-R1, R2, R3, Rm and R4) unchanged^20^. This deletion is embryonic lethal when homozygous (between embryonic day (E)15 and birth)^21^. In this model, *Nprl3*^-/-^ fetal liver demonstrates severe impairment of erythropoiesis compared to littermate controls. This is evident from visual inspection of embryos: healthy fetal liver at E13.5 is mostly comprised of haemoglobinising erythroblasts, whereas inability to fulfil this development in *Nprl3*^-/-^ embryos results in reduced fetal liver size (Fig. 1b), cellularity (Fig. 1c) and cell viability (Fig. 1d). Staging of erythroid differentiation based on Ter119/CD71 expression^22^ showed a profound block on terminal erythroid differentiation, with significantly fewer *Nprl3*^-/-^ erythroblasts reaching stage 3 (S3) (basophilic erythroblasts) compared to WT (Fig. 1e, Extended Data Fig. 1a and b). Notably, myelopoiesis and megakaryopoiesis are unimpaired in *Nprl3*^-/-^ fetal liver (Fig. 1f and g), showing an erythroid-specific dependence on sufficient *Nprl3* expression. Consistent with this erythroid specificity, common myeloid progenitors (CMPs) and megakaryocyte erythroid progenitors (MEPs) were both reduced in number in the *Nprl3*^-/-^ fetal liver, whereas granulocyte monocyte progenitors (GMPs) and c-kit^+^Sca-1^+^ haematopoietic stem and progenitor cells (HSPCs) were unchanged (measured by flow cytometry^23^; Extended Data. Fig. 1c). These results may suggest an early-stage bias away from erythroid development in addition to later-stage inhibition of erythropoiesis.

To assess the effects of *Nprl3*^-/-^ in adult erythropoiesis, independent of any potential systemic developmental defects, a fetal liver - bone marrow competitive chimaera model was established (Fig. 1h). One group of lethally irradiated CD45.1 *R26-dTomato* mice were adoptively transferred *Nprl3*^-/-^ CD45.2 fetal liver cells alongside *R26-dTomato* WT bone marrow cells, and another received WT CD45.2 fetal liver cells alongside *R26-dTomato* WT bone marrow cells. Equal numbers of fetal liver cells, and equal numbers of competitor bone marrow cells, were transferred between groups. As erythroid cells lose expression of CD45 antigens with maturity, the *R26-dTomato* ubiquitous reporter construct enabled the origin of mature RBCs to be defined in reconstituted mice as fetal liver (Tomato^-^) or bone marrow competitor (Tomato^+^).

Over 16 weeks of reconstitution, *Nprl3^-/-^* fetal liver-derived cells contributed poorly to the peripheral RBC population (Fig. 1i). After 16 weeks, we determined the contribution of *Nprl3^-/-^* cells through successive stages of bone marrow erythropoiesis, which showed a profound defect in terminal erythroid differentiation. *Nprl3^-/-^* cells comprised only 5-10% of basophilic, polychromatic and orthochromatic erythroblasts in the bone marrow, whereas WT fetal liver-derived cells comprised ∼60% (Fig. 1j; Extended Data. Fig. 1d; assessed by flow cytometry^24^). *Nprl3^-/-^* cells seemed to accumulate in the pro-erythroblast stage where their contribution was similar to WT fetal liver-derived cells (Extended Data. Fig. 1f), suggesting a differentiation block reminiscent of our observations in the fetal liver (Fig. 1e).

In addition to defects in terminal erythropoiesis in the fetal liver-bone marrow chimeras, we observed a competitive disadvantage at the earliest assessed haematopoietic bifurcation, with fewer MPPs formed by *Nprl3*^-/-^ fetal liver cells (Extended Data. Fig. 1f; haematopoietic progenitors were defined by flow cytometry using a previously characterised scheme^25^). This appeared to result in pan-haematopoietic disadvantage; *Nprl3^-/-^* fetal liver contributed comparatively poorly to the majority of progenitor types (Extended Data. Fig. 1f). This is not reflective of the comparatively normal numbers of non-erythroid cell types and their progenitors found in E13.5 *Nprl3*^-/-^ fetal liver (Fig. 1f and g). Notably, this HSC-level disadvantage was only observed in the competitive setting, where more ‘fit’ WT cells exist in the same haematopoietic system. The difference between these settings may also reflect that fetal liver erythropoiesis is driven by progenitors that develop independently of the HSC compartment, whereas adult bone marrow haematopoiesis is HSC-dependent^26^. Interestingly, HSCs and erythroid cells express the highest levels of *Nprl3* among haematopoietic cells in the mouse bone marrow (Extended Data. Fig. 1g) and it is known that HSC fitness is regulated by appropriate levels of mTORC1 signalling^27,28^.

Irrespective of any potential role for *Nprl3*^-/-^ in HSCs, the accumulation of *Nprl3^-/-^*pro-erythroblasts and subsequent defect in terminal erythropoiesis supports a role for *Nprl3*^-/-^ in erythroid differentiation, independent of systemic developmental defects in *Nprl3*^-/-^ embryos.

## Human *NPRL3*-KO erythroblasts respond defectively to a changing cellular environment

Homozygous *NPRL3* loss-of-function mutations have not been identified in humans, likely due to early lethality. To study NPRL3 in human erythropoiesis, we established an *in vitro* model of *NPRL3*-knockout (KO) in erythroid cells. High-efficiency *NPRL3*-KO was induced in primary human CD34+ HSPCs by genome-editing (Fig. 2a) and maintained throughout 15 days of erythroid-directed culture (Extended Data. Fig. 2a and b)^29,30^. To assess the ultimate erythroid productivity of single *NPRL3*-KO CFU-Es versus single negative control CFU-Es (NC; nucleofected with a non-targeting sgRNA), these cells were singly sorted into individual wells prior to erythroid culture. *NPRL3*-KO CFU-Es formed significantly fewer enucleated cells compared to their NC counterparts (Fig. 2b and Extended Data Fig. 2c), showing that loss of NPRL3 also impairs the output of human *in vitro* erythropoiesis. Monitoring of cell number during bulk cell culture indicated a proliferative burst in NC populations from day 7 of culture, resulting in a peak in cell number on day 9 (Extended Data. Fig. 2d). This was not observed in *NPRL3*-KO populations, suggesting a growth or differentiation impairment of *NPRL3*-KO erythroblasts. The culture is mostly comprised of basophilic erythroblasts at this time, and so this result is consistent with the loss of progression to S3 observed in murine *Nprl3^-/-^* fetal liver (Fig. 1e).

**Figure 2.**
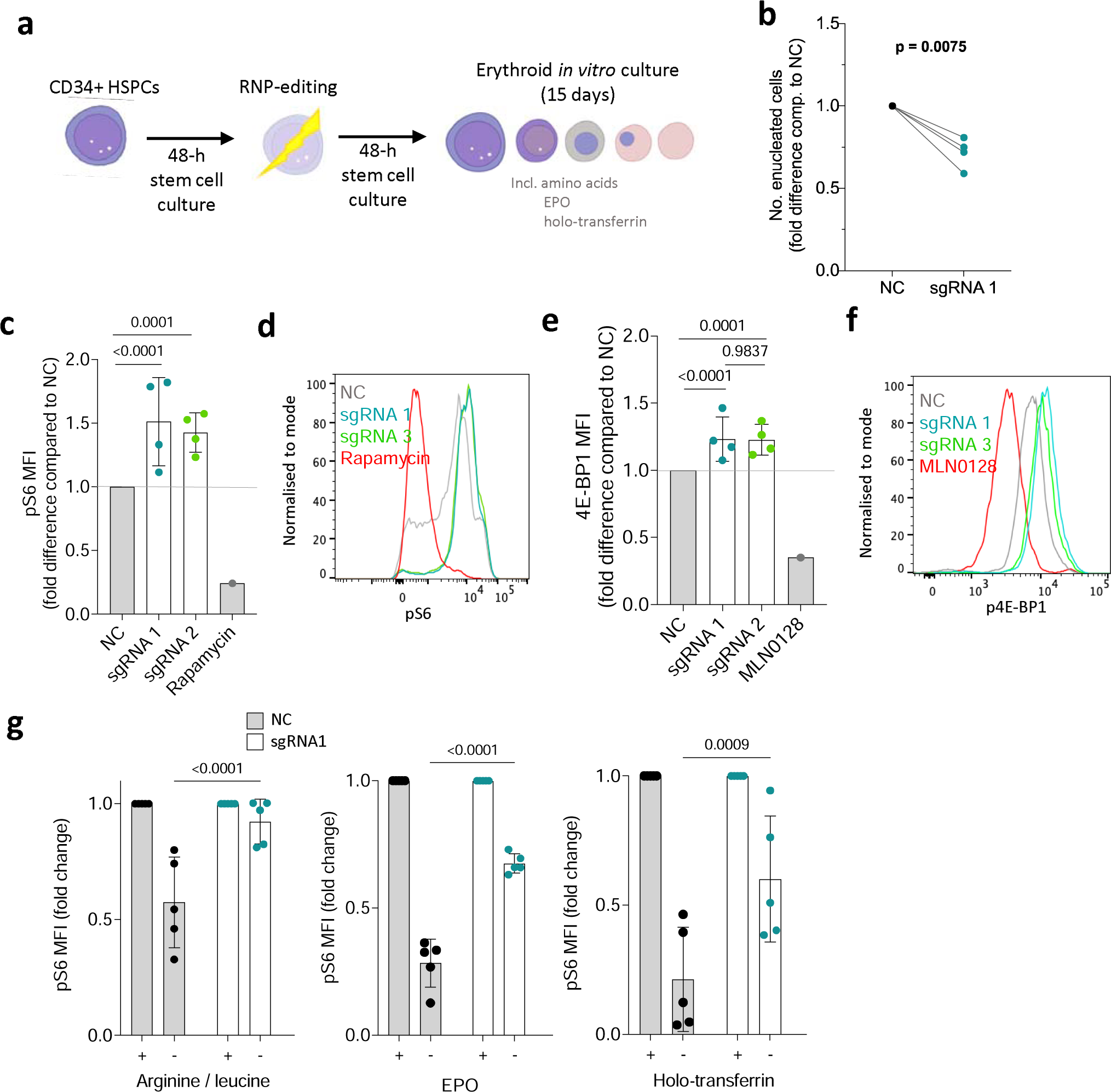
NPRL3-KO impairs primary human in vitro erythroid differentiation and erythroblast mTORC1 signalling. a) Schematic illustration: following extraction of human peripheral blood mononuclear cells, magnetic assisted cell sorting MACS was used to isolate CD34+ HSPCs. After a 48-h rest and expansion period, RNP-editing was performed to knockout NPRL3. After another rest period, the edited cells were exposed to an erythroid differentiation culture. b) Fold difference in the average number of enucleated cells formed by NPRL3-KO progenitors compared to NC by day 14. Four individual donors are indicated, and each pair of connected points represents the average from one individual, and each average is represented by the geometric mean. Analysed by one-tailed ratio paired t-test. c) pS6 median fluorescent intensity (MFI) measured 24 h post-media change by flow cytometry, also represented in d) using a histogram. e) p4E-BP1 MFI measured 24 h post-media change, also represented in f) using a histogram. Rapamycin and MLN0128 were included to indicate the dynamic scope of mTORC1 signalling measurable by pS6 MFI. Data are expressed as the mean ± SD. Fold reduction in pS6 MFI upon 4-h withdrawal of e) arginine and leucine, f) EPO and g) holo-transferrin, measured on day 11. Each point represents an individual donor. Analysed by Two-way ANOVA followed by Tukey’s test.

Human *NPRL3*-KO erythroblasts also showed overactive mTORC1 signalling, as measured by phosphoS6 (pS6; (Fig. 2c and d) and p4EB-P1 (Fig. 2e and f), close downstream effectors of mTORC1. This confirmed that NPRL3 deficiency results in increased mTORC1 activity by removing a means of negative mTORC1 regulation, demonstrating canonical GATOR-1 signalling in human erythroid cells for the first time. In cell lines and *Drosophila,* NPRL3 (as part of GATOR-1), inhibits mTORC1 activity in response to low amino acid availability, particularly arginine and leucine^31–33^. To test human erythroid cultures, edited erythroblasts were exposed to leucine and arginine withdrawal for 4 h before pS6 measurement. In such conditions, *NPRL3*-KO erythroblasts were unable to appropriately downregulate mTORC1 activity (Fig. 2f). Two key physiologic regulators of erythropoiesis are iron availability and erythropoietin (EPO). Interestingly, *NPRL3*-KO erythroblasts failed to appropriately regulate pS6 in response to iron deficiency and EPO deprivation (separately, in replete amino acid conditions; Fig. 2g and h). This shows that NPRL3 is required for appropriate integration of multiple stimuli that influence successful completion of erythropoiesis.

These results highlighted the importance of the NPRL3 - mTORC1 axis as a central metabolic regulator of erythropoiesis. We next revisited the fetal liver model to more closely assess changes in erythroid cell metabolism.

## Nprl3 maintains the metabolic profile of erythroid cells

Loss of Nprl3 increased erythroblast phospho-S6 (pS6) signalling in murine E13.5 *Nprl3*^-/-^ fetal liver (Fig. 3a), as in human erythroid cultures. Autophagy is an important contributor to cellular metabolism; it is required for erythropoiesis^34^ and is suppressed by activated mTORC1^18,35^. Therefore, we hypothesized impaired autophagy to be a possible outcome of overactive mTORC1 signalling induced by *Nprl3^-/-^*. Consistent with the role of Nprl3 as an mTORC1 inhibitor, autophagic flux was reduced in *Nprl3^-/-^* erythroblasts (S2) from E13.5 fetal liver compared to littermates (Fig. 3b). This likely contributes to their inability to efficiently mature to S3 (basophilic erythroblasts) and beyond.

**Figure 3.**
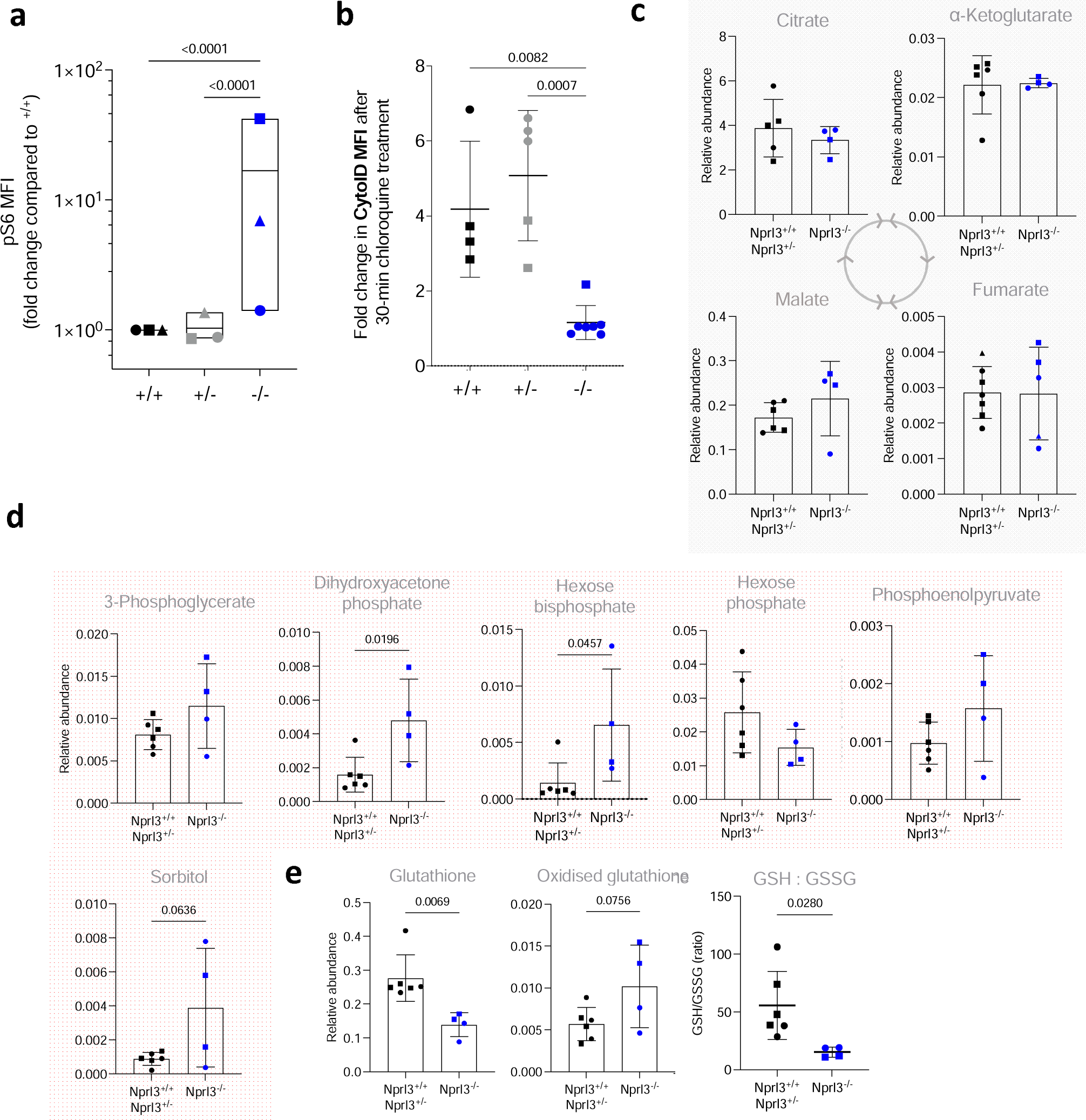
Metabolic effects of Nprl3^-/-^ in E13.5 fetal liver. a) pS6 MFI measured in S2 fetal liver erythroblasts (fold change compared to Nprl3^+/+^), following a 2-h 37°C incubation in RPMI with 5% FCS (otherwise un-supplemented). Each symbol indicates the average of the indicated genotype in one litter (n = 3 litters). Plotted on a log scale for visualisation. Analysed by Two-way ANOVA followed by Tukey’s test (performed on litter averages due to high baseline variability between experiments; this could be due to temporal instrumental variation of the flow cytometer used, or the speed of sample processing). b) Autophagic flux in S2 erythroblasts, as interpreted by difference in the MFI of Cyto-ID (an autophagosome dye) between untreated cells and cells treated with an inhibitor of autophagosome formation for 30 min. Each symbol indicates an embryo. Data are expressed as the mean ± SD. Analysed by One-way ANOVA followed by Tukey’s test. Relative abundance of c) glycolysis and d) Krebs cycle intermediates, in Nprl3^-/-^ Ter119+ cells versus littermates, measured by LC-MS. e) GSH:GSSG ratio, calculated from relative abundances of glutathione and oxidised glutathione. All LC-MS measurements were normalized to the abundance of myristic acid, employed as an internal control. All LC-MS data were compared by two-tailed t-test.

To further explore effects of *Nprl3*^-/-^ on metabolism, we analysed a panel of metabolic intermediates in Ter119+ erythroid cells from E13.5 fetal livers. Loss of Nprl3 did not alter concentrations of tricarboxylic acid cycle metabolites (Fig. 3c), amino acids or nucleotides (with increased levels of ATP relative to other nucleotides as expected in erythroid cells^36^) (Extended Data. Fig. 3). However, *Nprl3^-/-^* erythroid cells had increased abundance of hexose bisphosphate and dihydroxyacetone phosphate (DHAP; Fig. 3d), indicating relative dysregulation of glycolysis (a major mTORC1 target^37^) at the level of aldolase and triosephosphatase isomerase. DHAP is converted to glyceraldehyde-3 phosphate by triosephosphate isomerase (TPI), and genetic deficiency of TPI causes erythrocyte accumulation of DHAP and severe chronic haemolytic anaemia^38^. *Nprl3*^-/-^ was also associated with decreased glutathione (GSH) and an increased oxidised glutathione (GSSG) : GSH ratio (Fig. 3e), representing loss of reductive power and defective redox balance. Altogether, these specific changes in metabolites are consistent with altered mTORC1-glycolysis regulation in the Embden-Meyerhof pathway and decreased protection against oxidative damage. Since enzymopathies affecting either of these processes also cause erythrocytic defects^36^, these data are consistent with the erythroid deficiency in *Nprl3^-/-^*fetal livers and indicate that Nprl3-dependent control of metabolism is required for erythropoiesis.

## The α-globin enhancers increase erythroid *Nprl3* expression

Expression of *Nprl3* RNA increases with erythroid commitment^19^. We hypothesized that the α-globin enhancers drive this upregulation. To investigate this, we used transgenic breeding to eliminate *in cis* interaction between *Nprl3* and all murine α-globin enhancers. This involved the *Nprl3^+/-^* mouse (introduced in Fig. 1), where the transgenic allele has unperturbed α-globin enhancers^20^, but no *Nprl3* expression directed from its promoter. The second model harbours heterozygous deletion of all 5 α-globin enhancers (*Nprl3^+/AEKO^*; producing little or no α-globin RNA^39^), while *Nprl3* is unperturbed. When present independently, the *Nprl3^+/-^* and *Nprl3^+/AEKO^* genotypes do not affect the life expectancy or general health of adult mice. We gathered haematology readings from the peripheral blood of these mice and their littermates. *Nprl3^+/-^* adults demonstrate no difference in RBC count, size or Hb content compared to WT littermates (Extended Data Fig. 4a). *Nprl3^+/AEKO^* adults demonstrate normal RBC counts, with low MCV and Hb content (Extended Data Fig. 4b). As α-globin enhancers are required for α-globin expression on the allele *in cis*, *Nprl3^+/AEKO^* is equivalent to the expression of 2 of 4 functional α-globin genes (such as in α^0^-thalassaemia carriers). Such patients are often asymptomatic or have only mild anaemia, and their erythropoiesis is sufficient overall^40^.

Cross-breeding heterozygote *Nprl3^+/-^* and *Nprl3^+/AEKO^*mice enabled comparison of *Nprl3* RNA expression in the presence of full *Nprl3* expression (*Nprl3^+/+^*), heterozygous *Nprl3* (*Nprl3^+/-^*), homozygous *Nprl3* promoters with one allele lacking enhancers (*Nprl3^+/AEKO^*), and heterozygous *Nprl3* without possible enhancer influence (*Nprl3^AEKO/-^*). Importantly, all *in cis Nprl3*-α-globin enhancer interactions are eliminated in *Nprl3^AEKO/-^* animals. They express one allele with an intact *Nprl3* promoter and coding sequence, but lacking all α-globin enhancers, whereas the other allele lacks the *Nprl3* promoter, but all α-globin enhancers are expressed. These embryos were termed ‘*Nalph*’ (see schematic, Fig. 4a). Among 12 litters, *Nalph* was absent from 3 and was the least represented genotype overall (Extended Data. Fig. 4c), which may suggest a fitness disadvantage.

**Figure 4.**
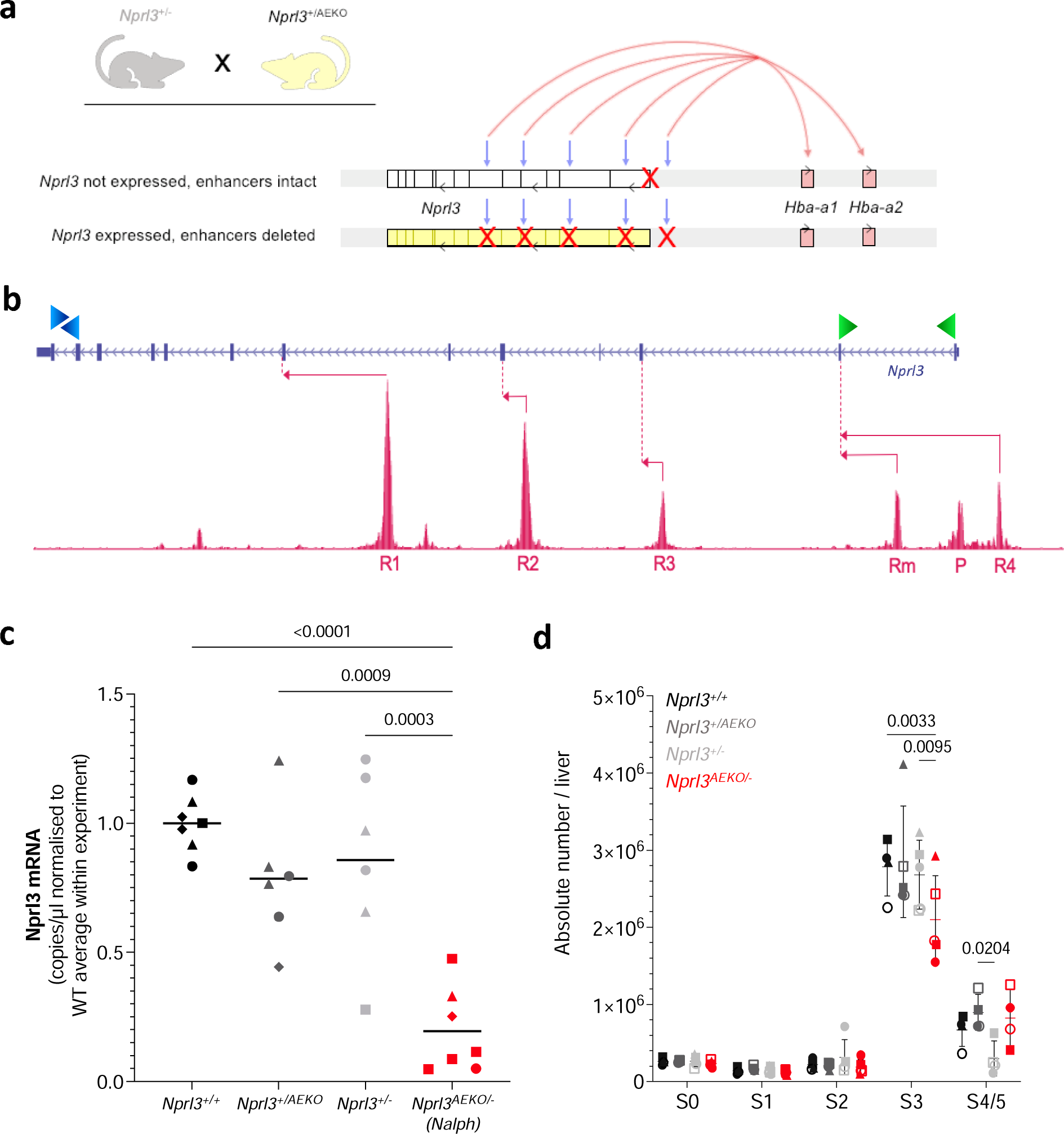
Elimination of interactions between the Nprl3 promoter and α-globin enhancers. a) Schematic illustrating the alleles inherited by the Nalph genotype, which is heterozygous for both the Nprl3 promoter deletion, and deletion of all α-globin enhancers. Grey arrows indicate the enhancer locations, and red arrows indicate enhancer induction of sufficient α-globin expression on one allele. b) Schematic illustrating the digital droplet assay design, with primer locations indicated by green (promoter-derived mRNA) and blue (mRNA + meRNA) arrowheads. Pink arrows indicate transcription starting from enhancers and their first splicing point. c) Promoter-specific Nprl3 mRNA expression in Nalph S3 erythroblasts compared to littermates (n = 4 litters). Analysed by One-way ANOVA followed by Tukey’s test. Each point represents an embryo, with litter represented by shape. d) Absolute number of erythroid cells in stages S0-S5 of differentiation in fetal livers of Nalph embryos vs. littermates (n = 5 litters). Data expressed as median with range. Analysed with by Two-way ANOVA followed by Tukey’s test.

To determine whether *Nprl3* expression is regulated by the α-globin enhancers in erythroid cells, we measured *Nprl3* mRNA in erythroid cells from all genotypes. *Nprl3* mRNA measurement has previously posed a challenge due to confounding measurement of full-length, spliced, polyadenylated enhancer-derived transcripts (meRNAs)^19^. We designed a high-sensitivity digital droplet PCR assay to measure promoter-specific *Nprl3* transcription (Fig. 4b). This was applied to cDNA collected from stage S3 sorted fetal liver cells. The levels of *Nprl3* RNA measured in *Nalph* erythroid cells were significantly lower than in littermates (Fig. 4c), and in particular, lower than when *Nprl3* lies *in cis* to functional α-globin enhancers in *Nprl3^+/-^*. Therefore, the α-globin enhancers regulate *Nprl3* in erythroid cells. This is supported by data from a high-resolution capture sequencing method (MCC-Seq)^20^, showing interaction between the *Nprl3* promoter and the R2 enhancer (Extended Data Fig. 4d). These findings are also consistent with Tri-C data showing that the *Nprl3* promoter is present with the α-globin genes and their enhancers in shared transcriptional hubs^11^.

To assess whether the enhancer-regulated transcriptional boost has a significant effect on the erythroid role of Nprl3, we compared the effects of enhanced and unenhanced levels of *Nprl3* RNA on erythroid development via flow cytometry on day E13.5. There was no difference in erythroid development between littermate *Nprl3^+/-^*, *Nprl3^+/AEKO^* and WT embryos (Fig. 4c). However, a lower number of erythroblasts in the S3 stage was observed of *Nalph* compared to littermates (Fig. 4d and Extended Data. Fig. 4e). The stage and direction of this effect is reminiscent of that observed in fully *Nprl3*-deficient (*Nprl3^-/-^*) fetal livers. The size of the difference is not as profound as in *Nprl3*^-/-^, consistent with decreased, but still functional, baseline *Nprl3* expression in *Nalph* cells that lack the erythroid transcriptional ‘boost’. These results indicate that the α-globin enhancers regulate *Nprl3* expression in erythroid cells, enabling optimal red blood cell production.

Overall, our data show that the locus containing *Nprl3*, the α-globin enhancers and α-globin controls developmental gene expression, allows interpretation of nutritional and signalling cues, and regulates erythroid cell metabolism, ultimately facilitating erythropoiesis.

## Discussion

Erythroid terminal differentiation requires intense anabolic activity to synthesize sufficient haemoglobin, as well as catabolism, both to recycle macromolecules and to enable the simplification from erythroblast to erythrocyte. We show that control of mTORC1 by Nprl3 is required during this process, and that erythropoiesis may be particularly vulnerable to Nprl3 deficiency.

We demonstrate that Nprl3 governs autophagic flux and regulates mTORC1 responses in erythroid cells not only to amino acid availability, but also to iron and EPO. It was recently found that cholesterol levels are also communicated to mTORC1 via GATOR-1 in HEK293T cells^41^. To our knowledge our results are the first demonstration in ‘non-transformed’ cells for a role of GATOR-1 (and NPRL3) acting as an integrator of environmental signals beyond amino acid availability and indicate a potential mechanism for the known regulation of mTORC1 by iron in erythroid cells^42^.

Metabolically, Nprl3 deficiency associates with dysregulation of redox control and the glycolytic Embden-Meyerhof pathways in erythroid cells. Erythrocytes are exposed to oxidative damage and require protective reductive power in the form of glutathione (GSH). Decreased GSH levels suggest that *Nprl3^-/-^*erythroid cells are relatively vulnerable to oxidative metabolic stressors. Erythroid proliferation and differentiation is supported by glycolysis^43^, and mTORC1 regulates glycolysis in mouse embryonic fibroblasts and cancerous cells via Hif-1 and c-myc^44,45^. Therefore, disruption of glycolysis due to mTORC1 overactivation could contribute to the *Nprl3^-/-^* erythroid differentiation defect. The increased DHAP level in *Nprl3^-/-^* erythroid cells is notable, both because DHAP accumulation also occurs in TPI deficiency (linked to severe anaemia)^36^, and because DHAP is known to activate mTORC1^46^. Therefore, mTORC1 overactivity driven by loss of Nprl3 may potentially be maintained and exacerbated by increased DHAP, further destabilizing metabolic control. Interestingly, glycolysis also supports HSC self-renewal^23^, and HSC commitment to the erythroid lineage^28^. If glycolytic dysregulation also occurs in *Nprl3^-/-^* HSCs this could contribute to their impairment in the competitive chimaera experiment (Extended Data. Fig. 1f) and represent a shared metabolic vulnerability of HSCs and erythroid cells.

We have shown that the α-globin enhancers upregulate *Nprl3* transcription in a manner important for erythropoiesis, independently of their critical role supporting α-globin expression. We hypothesize that the presence of the α-globin enhancers makes *Nprl3* expression more tuneable during erythropoiesis -helping to maintain sensitive regulation of mTORC1 signalling, particularly in response to environmental conditions and metabolic state. The phenomenon of enhancer sharing, in which lineage-specific enhancers regulate two different genes in the same locus, has been previously described. Shared enhancers regulate expression of the embryonic, fetal and adult globin genes at both the α-globin^47^ and β-globin loci^48,49^, and at the Hox clusters^50^. While these examples involve paralogous genes that act in the same pathway, enhancer sharing by otherwise unrelated genes has also been described (e.g. *Lnpk* in the HoxD cluster^51^). Interestingly, intronic enhancers within *Formin* critically control *Gremlin* expression in the developing limb, where *Formin* expression is dispensable^52,53^. Both *Formin* and *Gremlin* are involved in kidney development, though to the best of our knowledge, no enhancers have been demonstrated to co-regulate both genes in the kidney. We believe our work demonstrating co-regulation of *Nprl3* and the α-globin genes by the α-globin enhancers constitutes the first direct demonstration of functional cross-talk between distinct cellular pathways (mTORC1 and globin synthesis) relying on shared genomic enhancers, where each component is required for successful completion of a key biological process - in this case, erythropoiesis.

It is thought that, in a common ancestor of all vertebrates, an *Nprl3*-hosted regulatory element (corresponding to mammalian R1) provided a platform to regulate ancestral globin genes^4^. *Nprl3* has remained syntenic with the convergently evolving gnathostome haemoglobin α and agnathan monomeric haemoglobin^54,55^, both of which bind and transport oxygen. Together with the work presented here, this suggests that an ancient *Nprl3,* which regulated metabolism and possessed an enhancer, could have facilitated the tissue-specific expression and upregulation of early oxygen-carrying globins during the evolution of erythropoiesis.

Overall, in this study we show that Nprl3 is functionally important for metabolic control and development of erythroid cells, and that *Nprl3* expression is transcriptionally supported by the α-globin enhancers. We propose that the deep evolutionary coupling of α-globin and *Nprl3* has been maintained to allow this locus to control metabolism, as well as α-globin production, during erythropoiesis.

## Data availability

ATAC-Seq raw sequence data generated are available on the Gene Expression Omnibus (GEO) database under accession: GSE174110. Other data supporting the findings of this study are available within the paper and its Supplementary Information. Raw excel files are available upon request. No new code or large datasets were generated by this study.

## Supporting information

Flow gating

## Acknowledgements

JR BMS facility for indispensable technical assistance (Roo Bhasin, Jonathon Merril, Jordan Tanner). WIMM members for technical teaching and advice (Caroline Scott, Aude Anais, David Cruz Hernandez, Lucy Field, Lauren Murphy). The WIMM Flow Cytometry and Sequencing facilities. The VIB Metabolomics Core, Leuven Belgium. WIMM members for academic discussion (Tom Milne, Adam Wilkinson, Beth Psaila). Monika Kowalcyzk for creation of the *Nprl3* promoter-knockout mouse line and characterisation of meRNAs.

## Contributions

The present study was conceived by AP, HD and JF. Experimental design was carried out by, AP, MB, RN, MH, CB, JH, RB, MK. Experiments were performed by AP, JF, MT, SW, NW and AA. Data analysis was performed by AP, JF, JD. AP, JF, MK, NR, RN, DH and HD interpreted experimental results. AP and HD wrote the manuscript with contributions from all authors.

## Materials and Methods

### Mice

The constitutive *Nprl3*-promoter-knockout mouse model (*Nprl3^-/-^*; previously generated in the WIMM^20^) was rederived from cryopreserved sperm. The line was backcrossed onto a CB57BL/6J background by breeding heterozygotes with external WT CB57BL/6J mice (as monitored by the Transnetyx miniMUGA panel to >95% CB57BL/6J). The commercially available line, Rosa26^tdTomato^, was provided by the Mead group (WIMM). The background of these mice is C57BL6/J. The all-enhancer-knockout mouse was derived via the R2-only mouse line (generated by the Higgs group, WIMM^39,56^), by Ben Davies at the Wellcome Centre for Human Genetics (now Crick Institute, London). In brief, R2-only oocytes were microinjected with the following guide RNAs complexed with Cas9: R2 deletion right, GCCGTGACACTTCATGCTCA, and R2 deletion left, TACCTCCAAGGTTTTGCTC. These oocytes were reimplanted into female mice, whose consequent pups were screened by PCR to identify heterozygotes. This colony was crossed to C57BL6/J for two generations. Animal procedures were performed under the authority of UK Home Office project and personal licenses, in accordance with the Animals (Scientific Procedures) Act of 1998, and were approved by the University of Oxford ethical review committee. WT C57BL/6JOlaHsd mice used for breeding and backcrossing were ordered from Envigo Ltd. Mice were housed in individually ventilated cages and a standard diet (SDS Dietex Services; cat. no., 801161) was available *ad libitum*. Mice were euthanised in increasing concentrations of CO2, followed by confirmation through cervical dislocation. Timed matings were established for all fetal liver experiments. Mice were used between the ages off 6-18 weeks. Breeding pairs or trios were set up between 15:00 and 17:00, and plug checks were performed the next morning by 07:30. Observance of a vaginal plug marked embryonic day of development (E)0.5. After day E10.5, female mice which had displayed a plug could be visually checked to confirm pregnancy, before schedule 1 culling of the female and embryos on day E13.5. Fetal livers were collected by individual sterile dissection of embryos in cold PBS (0.5% BSA). Embryos used in the in-house transgenic cross model of *Nprl3^-/-^*and all-enhancers^+/-^ carried the following genotypes: *Nprl3^+/+^*, *Nprl3*^+/-^, *Nprl3^+/AEKO^* and *Nprl3^AEKO/-^* (Nalph), where ‘+’ indicates the WT Nprl3 allele, ‘-’ indicates the Npr3 promoter knockout, and ‘*^AEKO^’* indicates the all-enhancer knockout. The genotype of each parent varied between experiments.

### Fetal liver / bone marrow chimaeras

Female Rosa26-tdTomato mice (C57BL/6J; 12-18 weeks old) were lethally irradiated with 2x 4.5 Gys, delivered 4 h apart. Bulk fetal liver cells (WT or *Nprl3^-/-^*; C57BL/6J) and Rosa26-tdTomato competitor bone marrow cells were intravenously injected into the recipient tail 2-4 h after irradiation. R26-dTomato recipients were used to ensure that if complete lethal irradiation was not achieved, any residual progenitors could be grouped as ‘bone marrow competitors’. A total of 4.3x10^5^ fetal liver cells and 8.6x10^5^ tdTomato competitor bone marrow cells were injected per recipient (calculated for a 1:2 ratio, limited by the number of available *Nprl3^-/-^* FL cells). Recipients were administered with (0.16 mg/mL Enrofloxacin (Baytril), Bayer Corporation) in drinking water for 4 weeks post transplantation to prevent bacterial infection. Reconstitution of peripheral blood was assessed by flow cytometry, using 100 μl blood from the recipient tail vein at weeks 4, 8, 12 and 16.

### Flow cytometry for surface antigens

Tissues (bone marrow and fetal liver) were passed through 70- or 40-µm filters to create a single cell suspension. Cell suspensions were normalised for cell number, transferred to a 96-well round bottom plate, centrifuged at 1200 rpm for 5 min, and washed with 200 μl of PBS (2% FBS). Cells were incubated with anti-CD16/32 (FC receptor block) at 1:100 and LIVE/DEAD Fixable (1:400-1:800; cat no., 15519340; Invitrogen) in 20 μl PBS for 10 min. Fluorophore-conjugated antibodies were added at 2X listed dilutions and incubated for 15 min at 4 °C in the dark. Cells were washed in PBS (2% FBS) and analysed using an Invitrogen Attune or BD LSR Fortessa flow cytometer. Hoescht 33258 DNA stain (Invitrogen; cat. no., H3569) was added to human cell populations immediately prior to analysis cells in PBS (2% FBS) for viability assessment (human cells were not stained with LIVE/DEAD Fixable). See Supplementary material for antibody details.

### Flow cytometry for intracellular antigens

After processing as above and staining of surface antigens of interest, cells were fixed for 20 min in 150 μl 4% paraformaldehyde (BioLegend; cat. no., 420801) at 4 °C in darkness before a 30-min permeabilisation step (eBioscience; cat. no. 008333). Staining for intracellular markers was performed for 30 min in permeabilisation buffer, cells were then centrifuged at 1800 rpm for 2 min and resuspended in permeabilisation buffer for flow cytometric analysis.

### Autophagic flux analysis

A total of 3x10^5^ cells were suspended in RPMI + 5% FCS and plated (in duplicate per sample) in 250 μl in a 96-well plate triplicate. Chloroquine was added to one of the wells for each sample (50μM; Enzo Life Sciences; cat. no., ENZ-51031). The plate was incubated at 37 °C and 5% CO2 for 2 h. Cells were then processed using the CYTO-ID Autophagy Detection Kit (Enzo Life Sciences; cat. no., ENZ-51031), as per the manufacturer’s instructions. However, due to the small scale of this assay, only 125 μl diluted Cyto-ID dye was used per sample. Cells were then washed in the provided assay buffer, and surface antigen staining was performed as described above. Cells were immediately analysed by flow cytometry.

### Single cell sorting of HSPCs

A Sony MA900 Cell Sorter to deposit CFU-Es into single wells of round-bottomed 96-well plates containing pre-warmed HSPC media (see below).

### Isolation of primary CD34^+^ cells

Blood leukocyte cones were donated via the NHS Blood and Transplant centre (Oxford). Each cone was diluted 1:1 with PBS and layered onto Histopaque-1077 Hybri-Max (Sigma-Aldrich). The suspension was centrifuged at 1800rpm for 25 min at room temperature, with soft acceleration and deceleration. The interphase PBMC layer was collected with a pasteur pipette and washed twice with PBS. Cells were resuspended in MACS buffer (PBS, 0.5% BSA, 2 mM EDTA) for CD34^+^ magnetic selection using CD34 human MicroBeads (Miltenyi Biotec; cat. no., 130-046-702; 200 μl beads per cone, incubated at 4 °C on a roller for 30 min), a MidiMACS magnet (Miltenyi Biotec; cat. no., 130-042-302) and LS columns (Miltenyi Biotec; cat. no., 130-042-401). Cells were plated immediately into pre-warmed antibiotic-free HSPC media (StemSpan SFEM II; Stemcell Technologies; cat. no., 09605_C) supplemented with SCF (100 ng/ml; Peprotech; cat. no., 300-07), TPO (100 ng/ml; Peprotech; cat. no., 300-18) and Flt3 (100 ng/ml; Peprotech; cat. no., 300-19) at 0.25x106 cells/ml.

### *In vitro* human erythroid differentiation culture from CD34+ cells

This culture system has been previously established^29,30^. On day 0, HSPCs were seeded at 0.25x10^6^ cells/ml in phase 1 media (components and their concentrations are listed in Table 1). Cells were counted every 24-48 h from Day 3 using a NucleoCounter NC-3000 (Chemometec). Precisely, cell counts were performed on days 0 (post CD34+ isolation), 3, 4, 5, 7, 9 and 10. Until day 4, cell concentration was maintained at 0.25x10^5^ cells/ml. Until day 11, cell concentration was maintained at 5x10^5^ cells/ml after each cell count. On day 7, cells were resuspended in phase 2 media. On day 11, cells were resuspended in phase 3 media and maintained at 1x10^6^ cells/ml for the remainder of the culture. The NucleoCounter NC-3000 does not detect enucleated cells as live cells, therefore, as cell counts were not continued once from day 10. Enucleation rates are ∼20% by day 11, and ∼50% by day 14).

**Table 1.** *In vitro* liquid culture erythroid differentiation media constituents and their working concentrations.

| Component | Manufacturer | Catalog no. | Final concentration |  |
| --- | --- | --- | --- | --- |
| Iscoe's modified Dulbecco's media (IMDM) GlutaMAX | Gibco | 31980030 | - | 1,2,3 |
| Inactivated group AB Plasma | Dept. of Haematology, Oxford University Hospitals Trust | - | 3% | 1,2,3 |
| FBS | Sigma-Aldrich | - | 2% | 1,2,3 |
| Human holo- transferrin | Sigma-Aldrich | T0665 | 200µg/ml / 500µg/ml | 1,2/ 3 |
| Recombinant human insulin | Sigma-Aldrich | I9278 | 10µg/ml | 1,2,3 |
| Heparin sodium | Sigma-Aldrich | H3149 | 3U/ml | 1,2,3 |
| Recombinant erythropoietin- a | Bio-Rad | OBT1538 | 3U/ml | 1,2,3 |
| SCF | PeproTech | 300-07 | 10ng/ml | 1,2 |
| Penicillin/Streptomycin | Thermo Fisher Scientific | 15140122 | 100U/ml | 1,2,3 |
| Interleukin-3 | PeproTech | 200-03 | 1ng/ml | 1 |

To measure cell growth in terms of fold change in cell concentration over time, the concentration was normalised after every count from day 4 by. Day 7 represents replacement with phase 2 media. Each cell count was compared as a fold change to the preceding normalised concentration. As outlined above, for concentration normalisations, cell concentration was returned to the appropriate concentration for that media phase (0.25x10^6^ until day 4, 0.5x10^6^ until day 11 and 1x10^6^ thereafter).

For culture of singly-sorted HSPCs, cells were maintained in 150 μl phase 1 media in round-bottomed 96-well plates, within plastic boxes containing a reservoir of sterile water to avoid evaporative loses. Cells were not disturbed until day 7, at which point viable colonies were taken forward into phase 2, and media was changed as normal on day 11. The absolute number of enucleated cells was measured on day 11 using Hoescht 33342 (Invitrogen; cat. no., H3570; 0.5 μl 2.5 mg/ml Hoescht 33342 was added per well in 100 μl IMDM, and incubated at 37 °C for 25 min) for immediate analysis by flow cytometry (Hoescht 33342 not washed out).

### Ribonucleoprotein (RNP) editing

A total of 36-48 h post magnetic isolation, primary human CD34+ cells underwent RNP editing. Cas9 nuclease V3 (IDT; cat. no., 1081058) and sgRNAs (custom made by Synthego) were complexed at a 1:2.5 molarity ratio (Table 2). This was performed in PCR strip tubes with a 10-min incubation at 18 °C, then cells were kept on ice. Cells were washed in 1ml PBS in a V-bottom cryovial by centrifugation at 300g for 5 min, to aid complete removal of PBS before resuspension in P3 Nucleofection Solution (room temperature; P3 Primary Cell 4D-NucleofectorTM X Kit S; Lonza; cat. no., V4XP-3032). A total of 20 μl solution was used to resuspend 80,000 cells, and these cells were transferred to a cuvette (P3 Primary Cell 4D-NucleofectorTM X Kit S; Lonza) for nucleofection in a 4D-Nucleofector (Lonza) using the DZ100 programme. Cells were rested for 5 min before transferring into 100μl pre-warmed antibiotic-free HSPC media (supplemented as above). Edited cells were cultured for 24-48 h at 37 °C in 5% CO2 before differentiation was initiated.

**Table 2.**
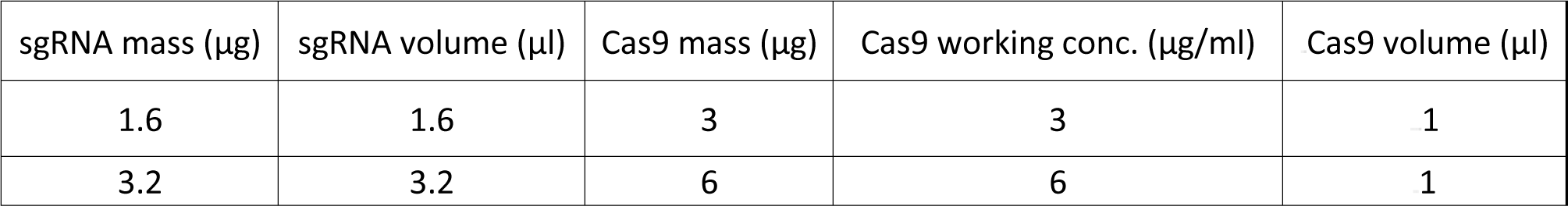
Example RNP complexing working conditions.

| sgRNA mass (μg) | sgRNA volume (μl) | Cas9 mass (μg) | Cas9 working conc. (μg/ml) | Cas9 volume (μl) |
| --- | --- | --- | --- | --- |
| 1.6 | 1.6 | 3 | 3 | 1 |
| 3.2 | 3.2 | 6 | 6 | 1 |

sgRNA sequences:

Nprl3 sgRNA 1: 5’-GCAACCAAGUCUGAAAUGUG-3’

Nprl3 sgRNA 3: 5’-UUGAUAAUGUGCGAUUUGUU-3’

Negative control sgRNA: non-editing sgRNA provided by Synthego (sequence not defined) not corresponding to any genomic region.

### Digital droplet PCR (ddPCR) RNA extraction

1-2x10^5^ sorted S3-stage fetal liver erythroblasts were centrifuged at 300 g for 5 min, the supernatant was removed and the cells were lysed in RLT+ buffer. The resulting lysate was used for RNA extraction using the RNA-easy Plus Mini Kit (Qiagen; cat.no., 74134) as per the manufacturer’s instructions.

### cDNA synthesis

Any contaminating DNA was removed from the RNA extract using the high efficiency TURBO DNA-free Kit (ThermoFisher Scientific; cat. no., AM1907), as per the manufacturer’s instructions. Using 2 μl RNA, cDNA was synthesised using the SuperScriptTM III First-Strand Synthesis System (Thermofisher Scientific; cat. no. 18080051).

### ddPCR reaction

The ddPCR reaction mix was prepared using 6 μl cDNA, ddPCR Supermix for Probes no dUTP (2X,; Bio-Rad; cat.no., 1863024), and contained final primer concentrations of 250 nM, and finl probe concentrations of 125 nM. The following primer and probe sequences were used, all custom made by Integrated DNA Technologies.

Primers and probe for *Nprl3* exon 1 - 2:

Exon1 - 2-Forward, 5’-CCATCAGCGTGATCCTGGTGAGCT-3’

Exon1 - 2-Reverse, 5’-CTGGTCATCAGCATGTTCGCCAGTG-3’

Exon1 - 2 probe: 5’-ACGCGGGGTGCTCCTGGCTCCTCTGGAA-3’ with 5’ HEX dye (PrimeTime®) and 3’ blackhole quencher (BHQ®-1).

Primers and probe for *Nprl3* exon 12 - 13:

Exon12 - 13-Forward, 5’-ACTCACCACTGAACAAGAGGATGACAG-3’

Exon12 - 13-Reverse, 5’-GCCGAGTGTTCTCATTGTACATGATCTC-3’

Exon1 - 2 probe: 5’-TGGTGGCGGCCACGGAAGTAGTGAAGGA-3’ with 5’ FAM dye (PrimeTime®) and 3’ blackhole quencher (BHQ®-1).

Primers for *Gapdh* housekeeping control:

Gapdh_ddPCR-Forward, 5’-AGGTCGGTGTGAACGGATTTG-3’

Gapdh_ddPCR-Reverse, 5’-TGTAGACCATGTAGTTGAGGTCA-3’

Probe for Gapdh housekeeping control: 5’-ATTGGGCGCCTGGTCACCAGGGCT-3’ with 5’ FAM dye (PrimeTime®) and 3’ blackhole quencher (BHQ®-1).

A foil plate lid was then applied at 180 °C for 3 sec. Droplets were generated using a BioRad Droplet Generator and Bio Rad ‘Oil for Probes’. The droplets were then exposed to the following thermocycling conditions to complete the ddPCR reaction:

1. Initial denaturation: 10 min 95 °C
2. Denaturation: 30 seconds at 94 °C
3. Annealing/Extension: 1 min at 60 °C (AEKD WT and KO) (Steps 2 and repeated for 40 cycles)
4. Enzyme inactivation: 10 min at 98 °C

(Ramp rate of 2 °C/sec applied at all stages). Product was measured using a QX200 Droplet Reader (Bio-Rad).

### Analysis

The presented mRNA/eRNA readings (copies/µl) are generated by the QX200 Droplet Reader, using an internal BioRad internal algorithm. To precisely normalise for input concentration, copies/µl readings were first normalised to that of a reference gene, Gapdh. Next, they were normalised to the average of readings for the control genotype (*Nprl3^+/+^ AEKO^+/+^*) in each litter.

### Metabolomics

Ter119+ cells were extracted using Anti-Ter119 MicroBeads (Milenyi Biotec; cat. no., 130-049-901), and LS columns (Milenyi Biotec), according to the manufacturer’s instructions. All reagents were kept on ice. Ter119+ cells were washed in PBS, pelleted, and immediately exposed to 80% methanol containing 2 µM d27 myristic acid, aand incubated for 3 min on ice. Samples were centrifuged at 17,000 g for 15 min. The supernatant was transferred to a new Eppendorf, and stored at -80°C until metabolite analysis.

Mass Spectrometry measurements were performed using Dionex UltiMate 3000 LC System (Thermo Scientific) coupled to a Q Exactive Orbitrap mass spectrometer (Thermo Scientific) operated in negative mode. 10 μl sample was injected onto a Poroshell 120 HILIC-Z PEEK Column (Agilent InfinityLab). A linear gradient was carried out starting with 90% solvent A (acetonitrile) and 10% solvent B (10 mM Na-acetate in mqH2O, pH 9.3). From 2 to 12 min the gradient changed to 60% B. The gradient was kept on 60% B for 3 minutes and followed by a decrease to 10% B. The chromatography was stopped at 25 min. The flow was kept constant at 0.25 ml/min. The column temperature was kept constant at 25°C. The mass spectrometer operated in full scan (range [70.0000-1050.0000]) and negative mode using a spray voltage of 2.8 kV, capillary temperature of 320°C, sheath gas at 45, auxiliary gas at 10. AGC target was set at 3.0E+006 using a resolution of 70000. Data collection was performed using the Xcalibur software (Thermo Scientific). The data analyses were performed by integrating the peak areas (El-Maven – Polly - Elucidata).

Absolute abundance measurements were normalised to the internal standard (myristic acid or D7-glucose) to generate relative abundance.

### Statistics

Statistics were performed in GraphPad Prism software (Version 9.4.1). Statistical significance was defined to be indicated by p < 0.05. Normality checks were performed before parametric testing. T-tests were performed when comparing the means of two groups, and paired if samples were matched between groups. ANOVA was used to make multiple comparisons between >2 normally distributed groups. One-way or Two-way ANOVA was performed depending on the number of independent variables (one or two, respectively). For optimal power when multiple comparisons testing, the post-hoc test used depended on the comparisons being made. Tukey’s test was performed when comparing every row (or column) mean with every other row (or column) mean. Šídák’s test was used to provide more power when comparing rows (or columns) within columns (or rows). When including individual repeat datapoints in grouped statistical analysis, a consistent number of repeats for every group is required to perform ANOVA. In the rare situation that single repeat measurements were missing for particular groups, a Prism Mixed-effects model was performed to maintain the power of using replicate data, and to generate comparisons equivalent to those made by ANOVA. For the - squared test, observed and expected numbers were compared with 3 degrees of freedom.

### Antibodies

**Table 3.**
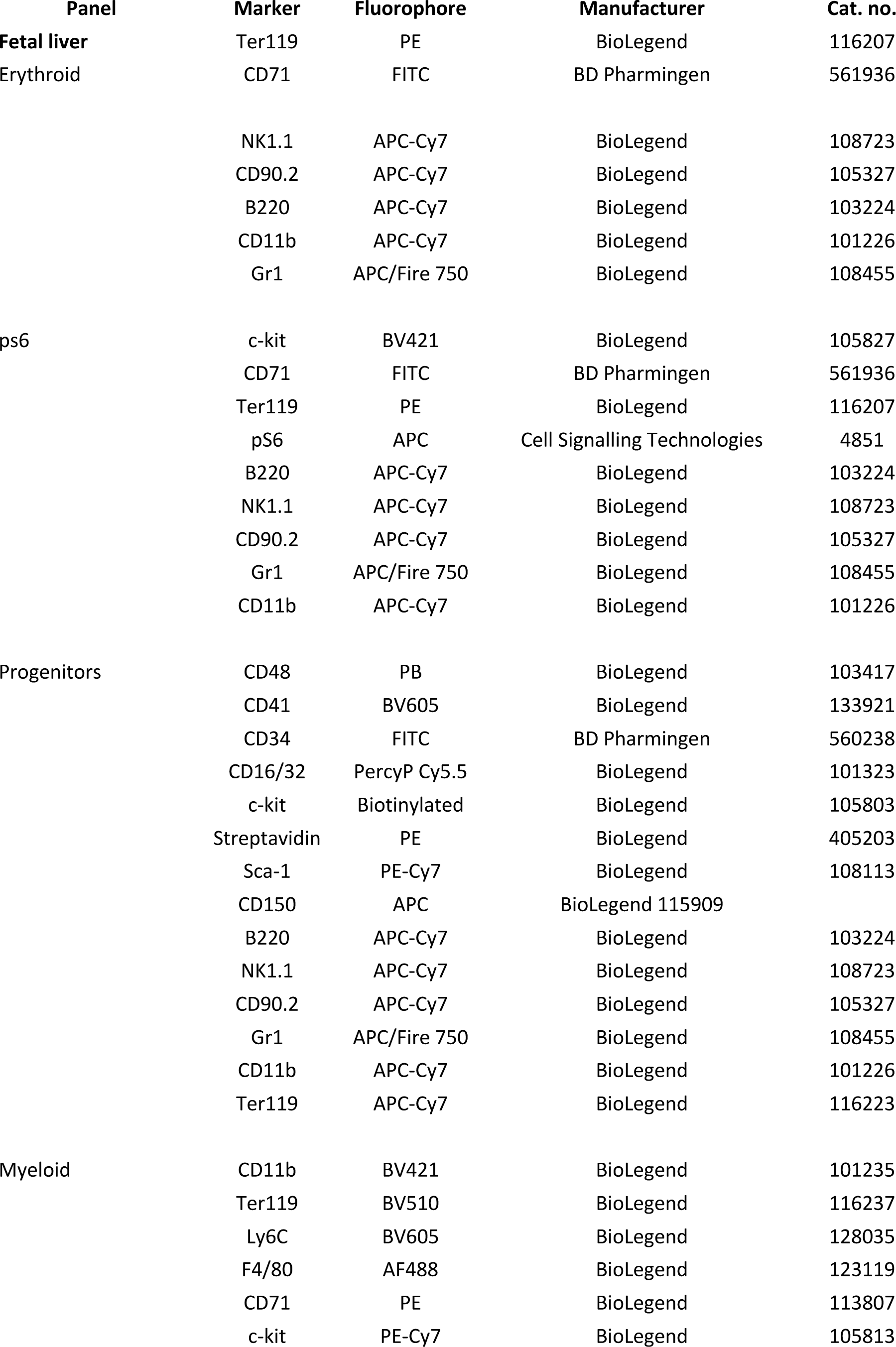
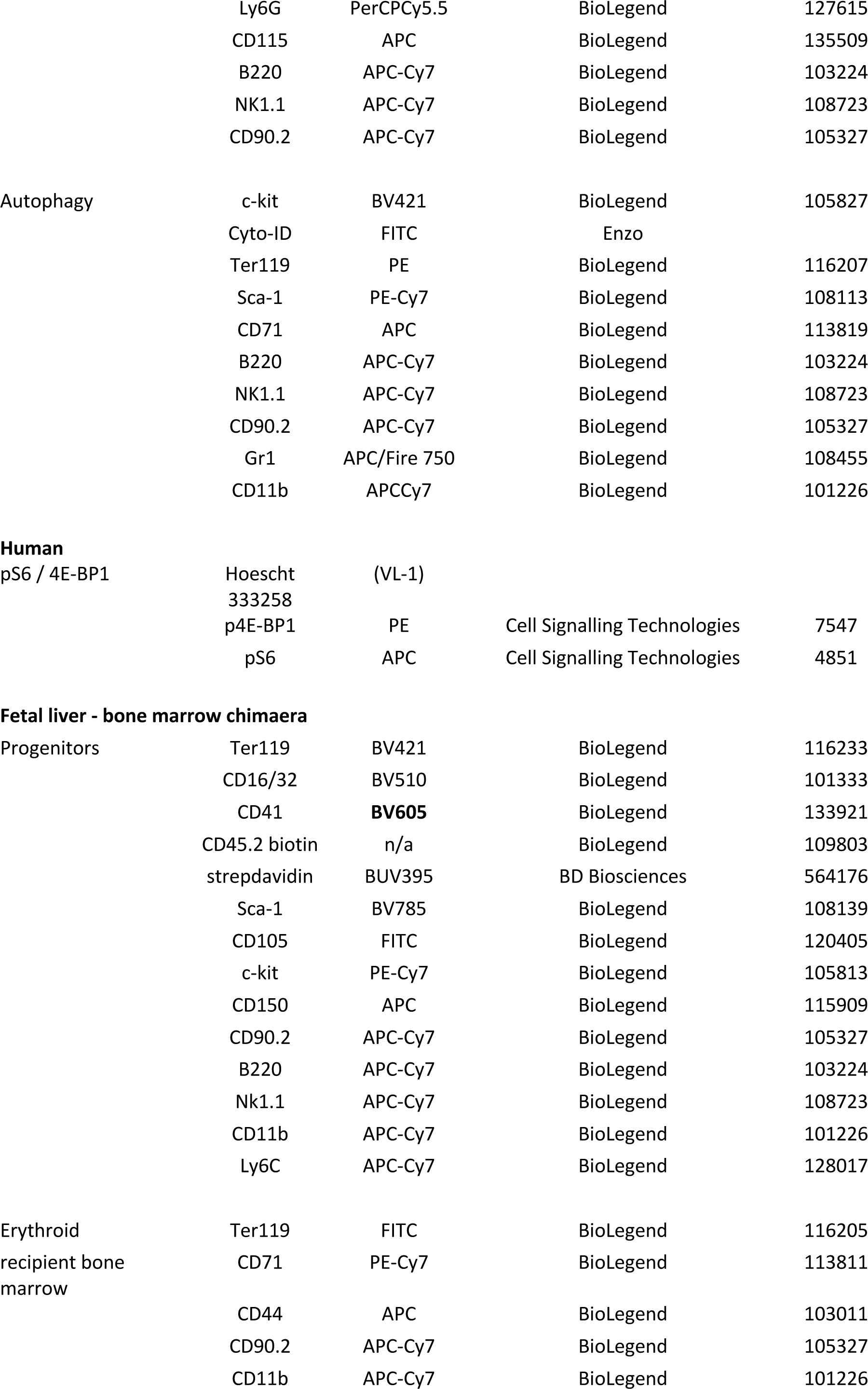
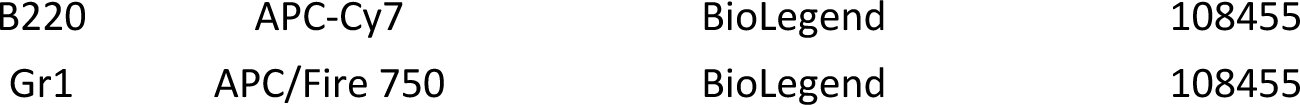
Flow cytometry antibodies used in this study, grouped according to experimental panel.

---

**Extended Data. Fig. 1.**
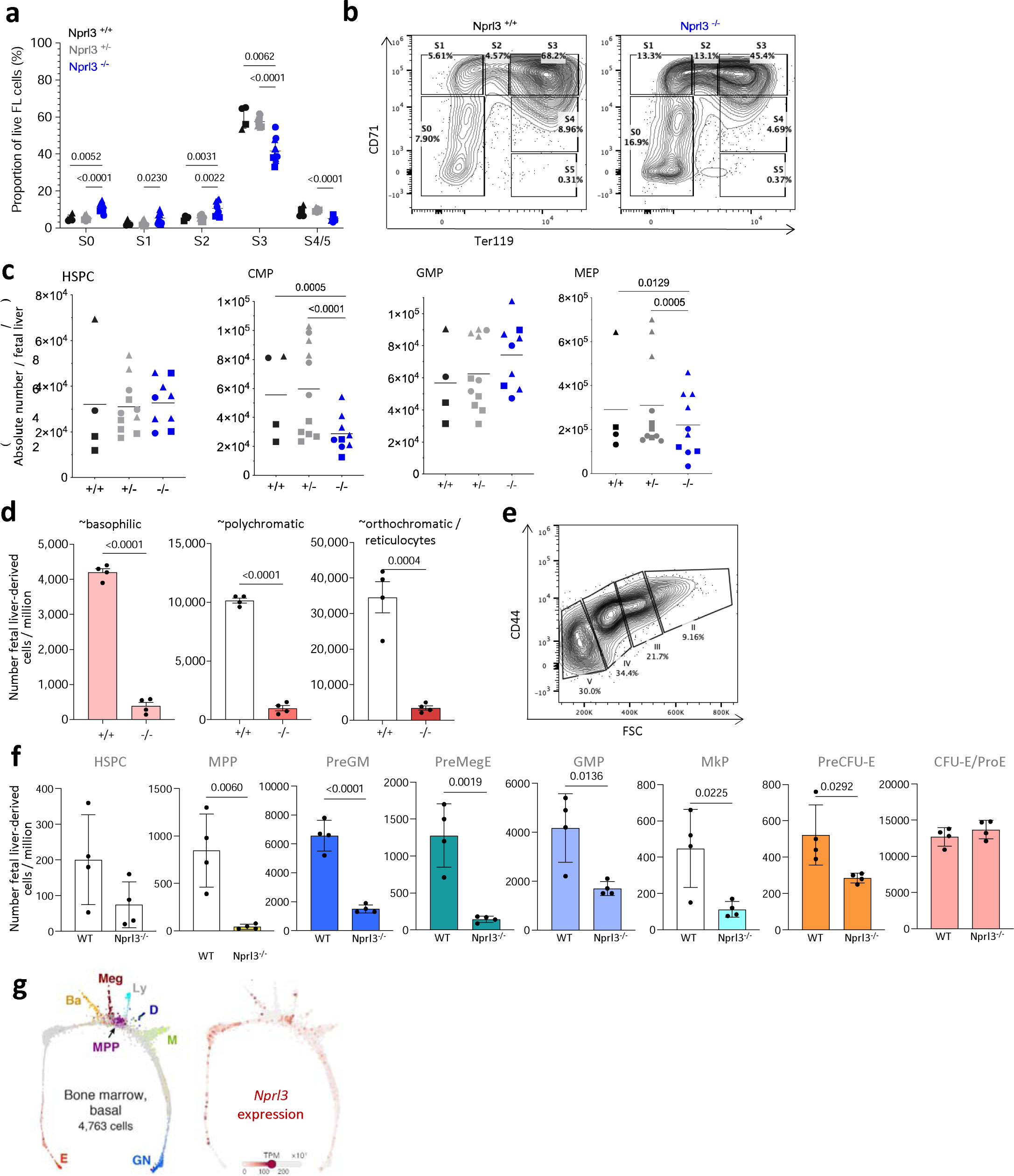
Effects of Nprl3^-/-^ in the fetal liver and in competitive chimaeric mice. a) Erythroid stages as a proportion of total live fetal liver cells according to genotype (n = 3 litters). Points indicate individual embryos. Data are expressed as the mean ± SD. Analysed by Two-way ANOVA followed by Tukey’s test. b) Example gating of erythroid stages in the fetal liver at E13.5. c) Absolute number per fetal liver of: total c-kit+ cells of lineage-Sca-1+ HSPCs, CD34+ CD16/32+ GMPs, CD34+ CD16/32-CMPs and CD34-CD16/32-MEPs (n = 5 litters). Each point represents an individual embryo, and litter indicated by shape of point. Data are expressed as the mean ± SD, and analysed by Two-way ANOVA (grouped by litter) followed by Tukey’s test. d) Absolute number of erythroblasts in stages II, II and IV derived from either WT or Nprl3^-/-^ fetal liver cells, per million recipient bone marrow cells. Analysed in recipient bone marrow at week 16 post-transplantation (n = 8 mice per group). Data analysed by two-tailed t-test, and expressed as the mean ± SD. e) Example gating of erythroid stages in the bone marrow. f) Absolute number of fetal liver-derived (CD45.2+ Tomato-) progenitors per million recipient bone marrow cells at week 16 post transplantation (n = 4 mice per group). Data analysed by two-tailed t-test, and expressed as the mean ± SD. g) Single cell RNA-Seq measured from mouse bone marrow-derived cells. Adapted from an online tool published in Tusi et al. 2018. https://kleintools.hms.harvard.edu/paper_websites/tusi_et_al/.

**Extended Data. Fig. 2.**
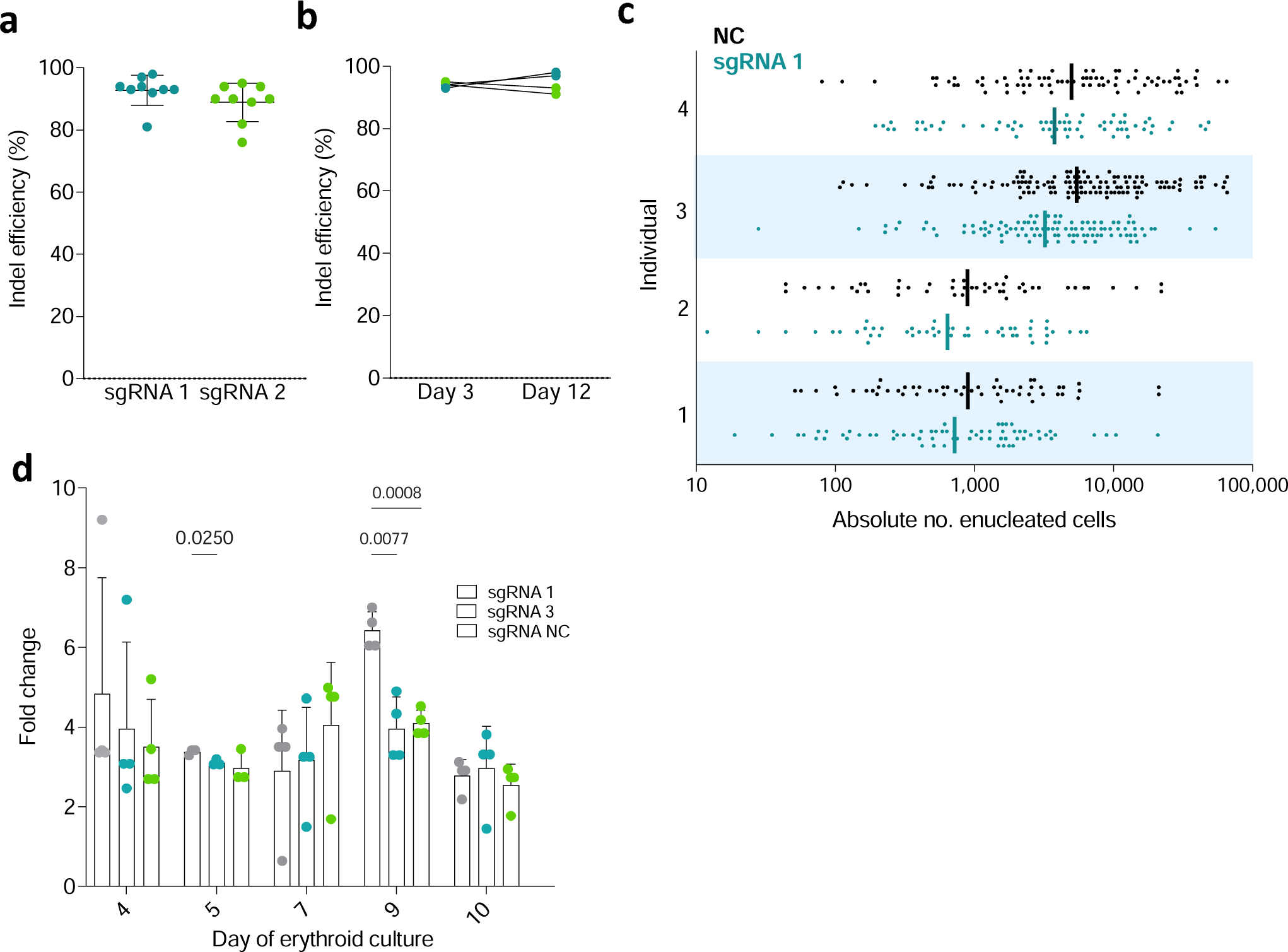
Effects of NPRL3-KO on primary human in vitro erythroid differentiation. a) Editing efficiency (% reads with identical knockout indel) achieved with NPRL3-targeting sgRNAs, sgRNA 1 and sgRNA 3. Each point represents a nucleofected donor population. b) Maintenance of editing efficiency from day 3 to day 12 of erythroid culture. c) Absolute number of enucleated cells formed from a single starting cell (each point represents a single starting cell). Bars indicate the geometric mean. Four individual donors are represented, stacked. d) Fold change in cell concentration throughout erythroid differentiation compared to previous dilution. Cells were counted on days 3, 4, 5, 7, 9 and 10, and concentration was normalised after every count from day 4 by media top up (except for day 7, which represents replacement with phase 2 media). Plotted points indicate the fold change in cell concentration relative to the normalised concentration established after the previous count point. Data represents 3 individual donors, expressed as the mean ± SD, and analysed by Prism Mixed-effects analysis matched by donor individual, followed by Tukey’s test.

**Extended Data Fig. 3.**
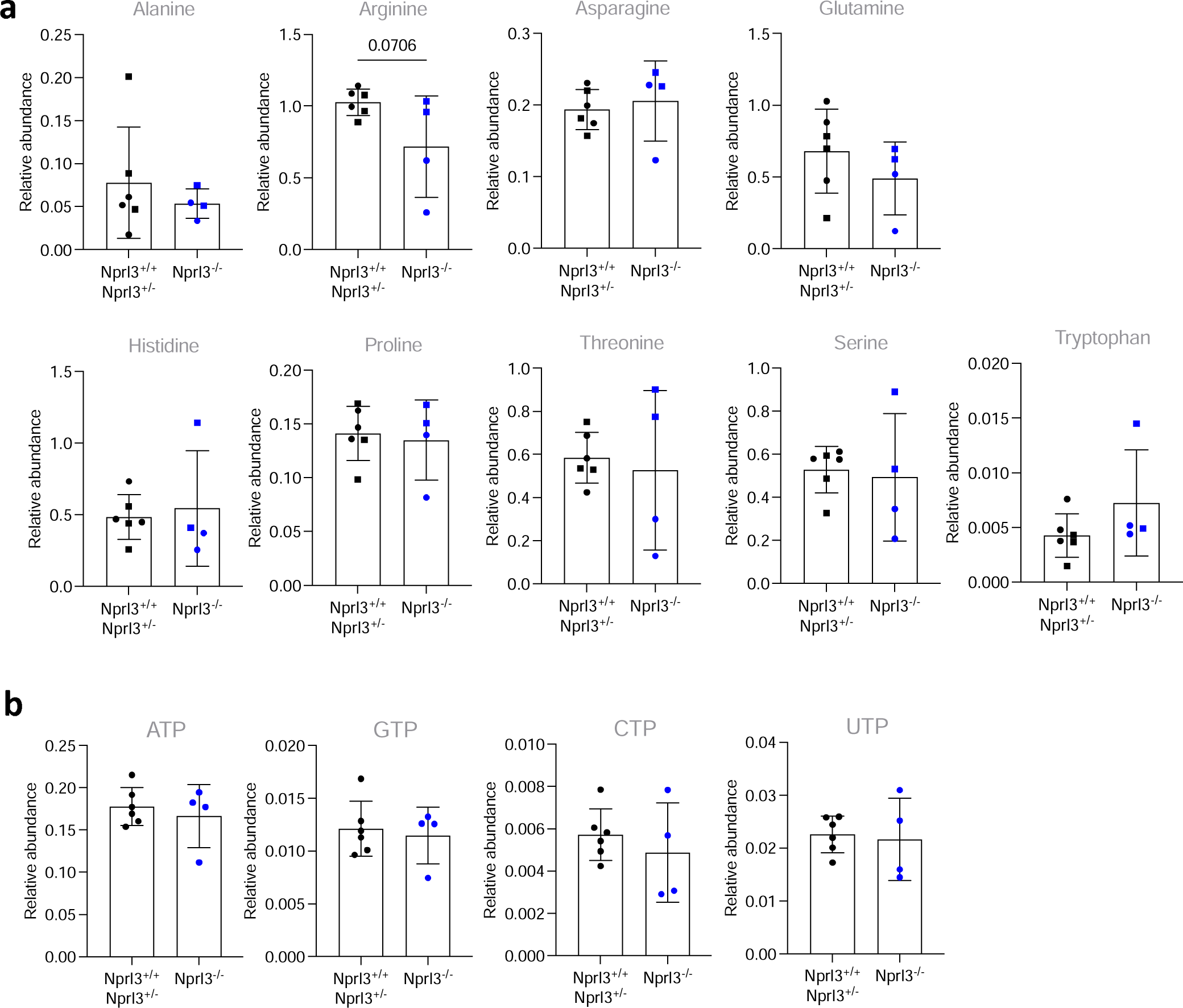
Amino acids and nucleotides measured by mass spectrometry. a) Relative abundance of amino acids, measured by LC-MS, normalised to D7-glucose (internal control). B) Relative abundance of nucleotides, measured by LC-MS normalised to myristic acid (internal control). All LC-MS data were compared by two-tailed t-test.

**Extended Data. Fig. 4.**
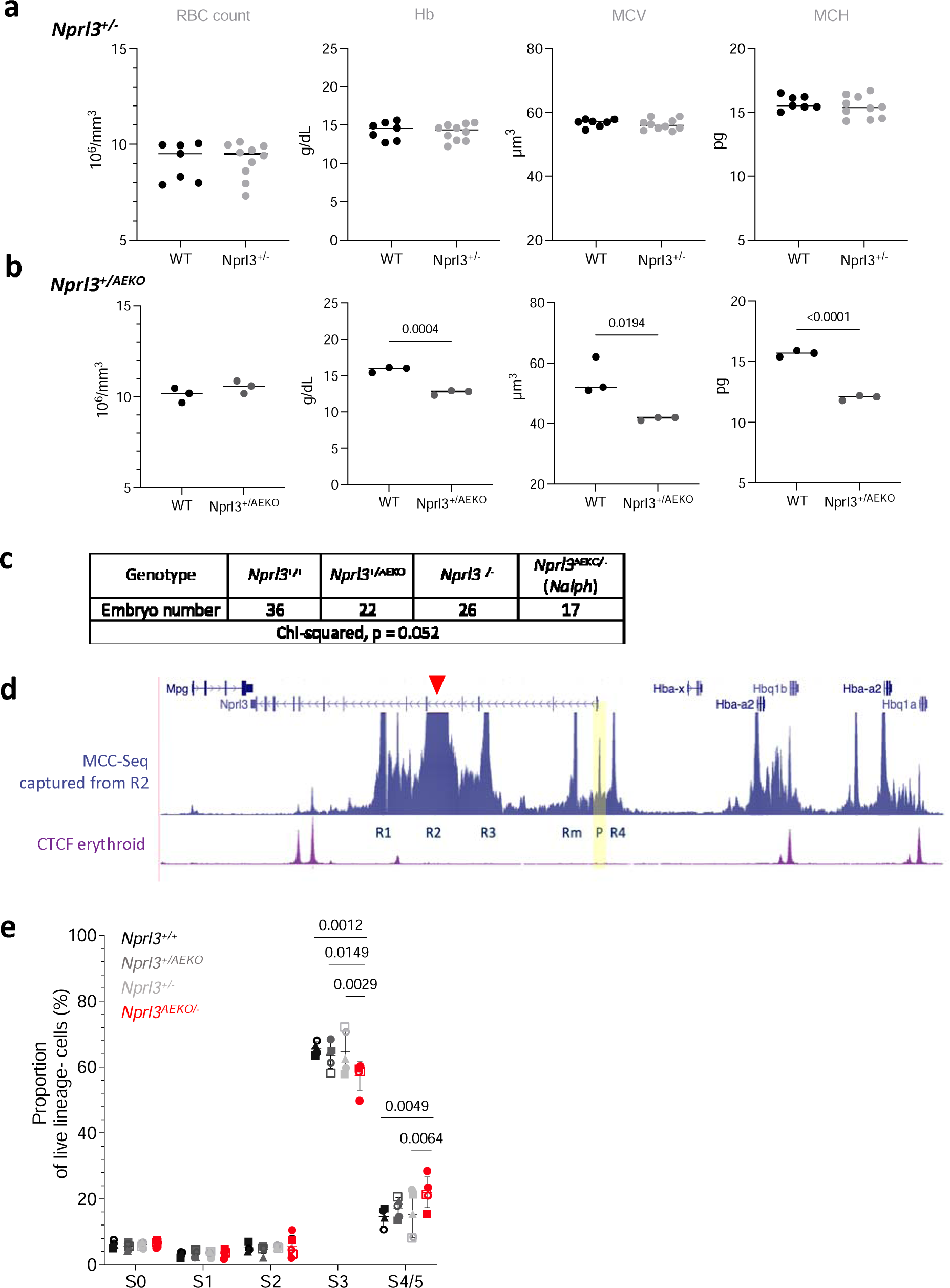
Further characterisation of the Nprl3-α-globin transcriptional hub. a, b) Peripheral blood measurements (RBC count, Haemoglobin (Hb), mean corpuscular volume (MCV) and mean corpuscular haemoglobin (MCH)) of a) Nprl3^+/-^, and b) Nprl3^+/AEKO^ adults and their littermate WT controls. Analysed by two-tailed t-test. Data expressed as the mean ± SD. c) Number of embryos of each genotype across 12 litters of Nprl3^+/-^ and Nprl3^+/AEKO^ animal crosses. d) MCC-Seq snapshot captured from the R2 enhancer (anchor point indicated by red arrowhead). Yellow panel highlights the interaction of the R1 enhancer with the Nprl3 promoter (P). e) Proportions of erythroid cells in stages S0-S5 of differentiation in fetal livers of Nalph embryos vs. littermates (n = 5 litters). Data are expressed as the mean ± SD. Analysed by Prism Mixed-effects followed by Tukey’s test. Each point represents an embryo, with litter represented by shape.

