## Supplementary material for "Ancient genomic linkage couples metabolism with erythroid development": Flow gating

### Slide 1
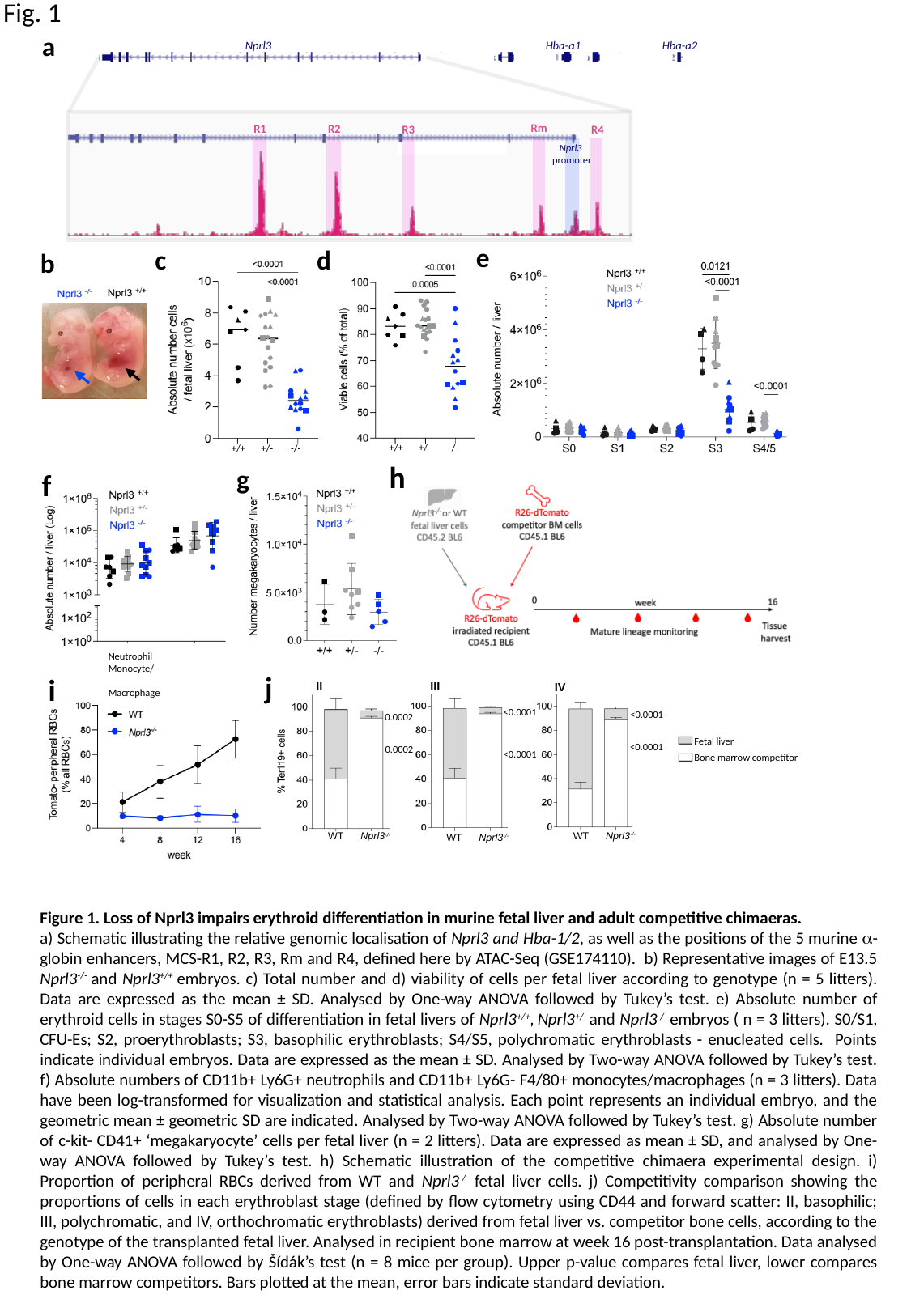

Fig. 1
a
Nprl3
Hba-a2
Hba-a1
Rm
R1
R2
R3
R4
Nprl3
promoter
e
c
d
b
+/+ +/- -/-
+/+ +/- -/-
h
g
f
Neutrophil	Monocyte/
	Macrophage
j
i
III
II
IV
<0.0001
0.0002
<0.0001
<0.0001
<0.0001
0.0002
WT Nprl3^-/-^
WT Nprl3^-/-^
WT Nprl3^-/-^
Fetal liver
Bone marrow competitor
Figure 1. Loss of Nprl3 impairs erythroid differentiation in murine fetal liver and adult competitive chimaeras.
a) Schematic illustrating the relative genomic localisation of Nprl3 and Hba-1/2, as well as the positions of the 5 murine α-globin enhancers, MCS-R1, R2, R3, Rm and R4, defined here by ATAC-Seq (GSE174110). b) Representative images of E13.5 Nprl3^-/-^ and Nprl3^+/+^ embryos. c) Total number and d) viability of cells per fetal liver according to genotype (n = 5 litters). Data are expressed as the mean ± SD. Analysed by One-way ANOVA followed by Tukey’s test. e) Absolute number of erythroid cells in stages S0-S5 of differentiation in fetal livers of Nprl3^+/+^, Nprl3^+/-^ and Nprl3^-/-^ embryos ( n = 3 litters). S0/S1, CFU-Es; S2, proerythroblasts; S3, basophilic erythroblasts; S4/S5, polychromatic erythroblasts - enucleated cells. Points indicate individual embryos. Data are expressed as the mean ± SD. Analysed by Two-way ANOVA followed by Tukey’s test. f) Absolute numbers of CD11b+ Ly6G+ neutrophils and CD11b+ Ly6G- F4/80+ monocytes/macrophages (n = 3 litters). Data have been log-transformed for visualization and statistical analysis. Each point represents an individual embryo, and the geometric mean ± geometric SD are indicated. Analysed by Two-way ANOVA followed by Tukey’s test. g) Absolute number of c-kit- CD41+ ‘megakaryocyte’ cells per fetal liver (n = 2 litters). Data are expressed as mean ± SD, and analysed by One-way ANOVA followed by Tukey’s test. h) Schematic illustration of the competitive chimaera experimental design. i) Proportion of peripheral RBCs derived from WT and Nprl3^-/-^ fetal liver cells. j) Competitivity comparison showing the proportions of cells in each erythroblast stage (defined by flow cytometry using CD44 and forward scatter: II, basophilic; III, polychromatic, and IV, orthochromatic erythroblasts) derived from fetal liver vs. competitor bone cells, according to the genotype of the transplanted fetal liver. Analysed in recipient bone marrow at week 16 post-transplantation. Data analysed by One-way ANOVA followed by Šídák’s test (n = 8 mice per group). Upper p-value compares fetal liver, lower compares bone marrow competitors. Bars plotted at the mean, error bars indicate standard deviation.

### Slide 2
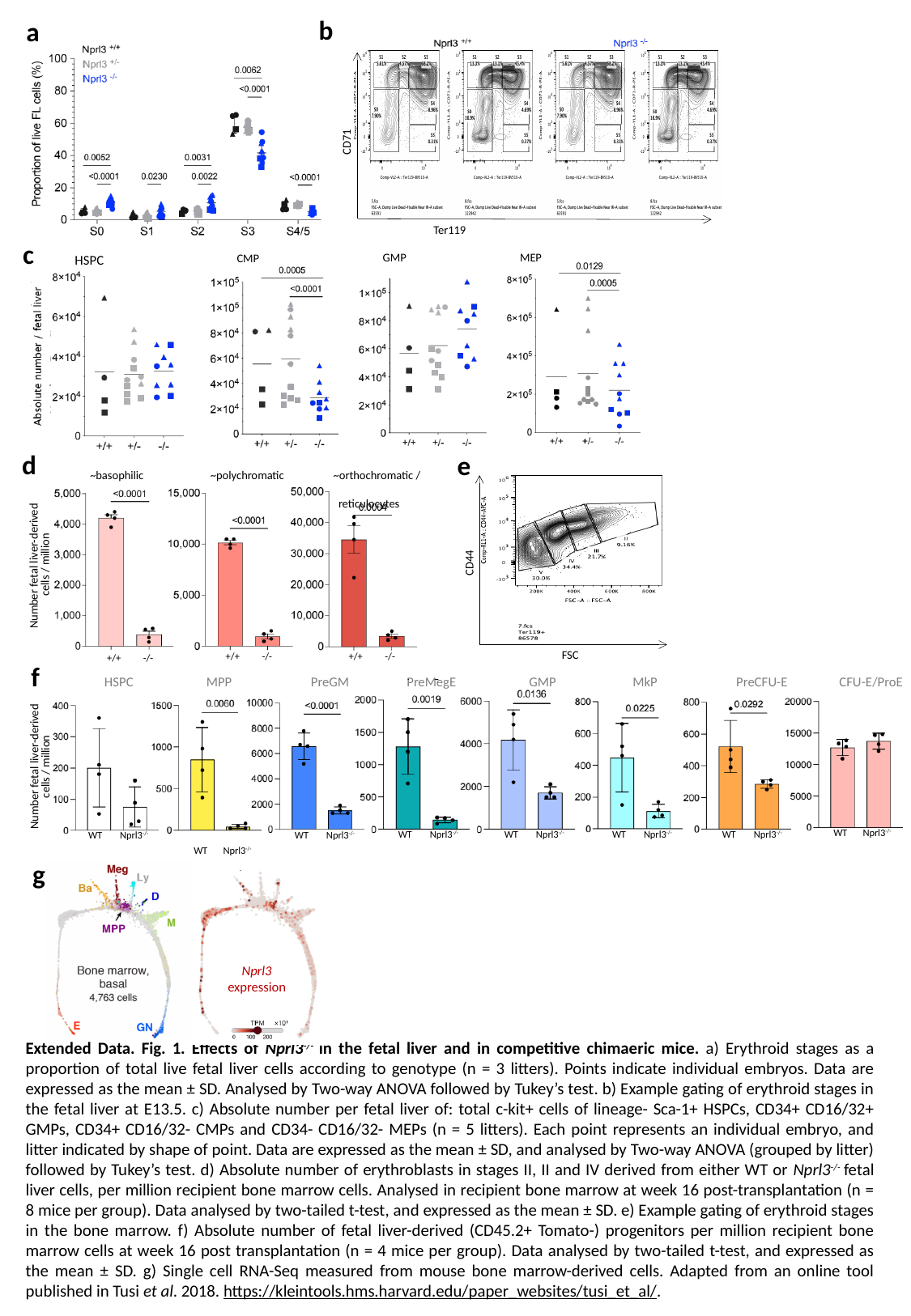

b
a
CD71
Ter119
c
GMP
MEP
CMP
HSPC
d
e
~polychromatic
~orthochromatic /
 reticulocytes
~basophilic
+/+ -/-
+/+ -/-
+/+ -/-
Number fetal liver-derived
cells / million
CD44
FSC-A
FSC
f
 HSPC MPP PreGM PreMegE GMP	 MkP PreCFU-E CFU-E/ProE
Number fetal liver-derived
cells / million
 WT Nprl3^-/-^
WT Nprl3^-/-^
WT Nprl3^-/-^
WT Nprl3^-/-^
WT Nprl3^-/-^
WT Nprl3^-/-^
WT Nprl3^-/-^
 WT Nprl3^-/-^
g
Nprl3
expression
Extended Data. Fig. 1. Effects of Nprl3^-/-^ in the fetal liver and in competitive chimaeric mice. a) Erythroid stages as a proportion of total live fetal liver cells according to genotype (n = 3 litters). Points indicate individual embryos. Data are expressed as the mean ± SD. Analysed by Two-way ANOVA followed by Tukey’s test. b) Example gating of erythroid stages in the fetal liver at E13.5. c) Absolute number per fetal liver of: total c-kit+ cells of lineage- Sca-1+ HSPCs, CD34+ CD16/32+ GMPs, CD34+ CD16/32- CMPs and CD34- CD16/32- MEPs (n = 5 litters). Each point represents an individual embryo, and litter indicated by shape of point. Data are expressed as the mean ± SD, and analysed by Two-way ANOVA (grouped by litter) followed by Tukey’s test. d) Absolute number of erythroblasts in stages II, II and IV derived from either WT or Nprl3^-/-^ fetal liver cells, per million recipient bone marrow cells. Analysed in recipient bone marrow at week 16 post-transplantation (n = 8 mice per group). Data analysed by two-tailed t-test, and expressed as the mean ± SD. e) Example gating of erythroid stages in the bone marrow. f) Absolute number of fetal liver-derived (CD45.2+ Tomato-) progenitors per million recipient bone marrow cells at week 16 post transplantation (n = 4 mice per group). Data analysed by two-tailed t-test, and expressed as the mean ± SD. g) Single cell RNA-Seq measured from mouse bone marrow-derived cells. Adapted from an online tool published in Tusi et al. 2018. https://kleintools.hms.harvard.edu/paper_websites/tusi_et_al/.

### Slide 3
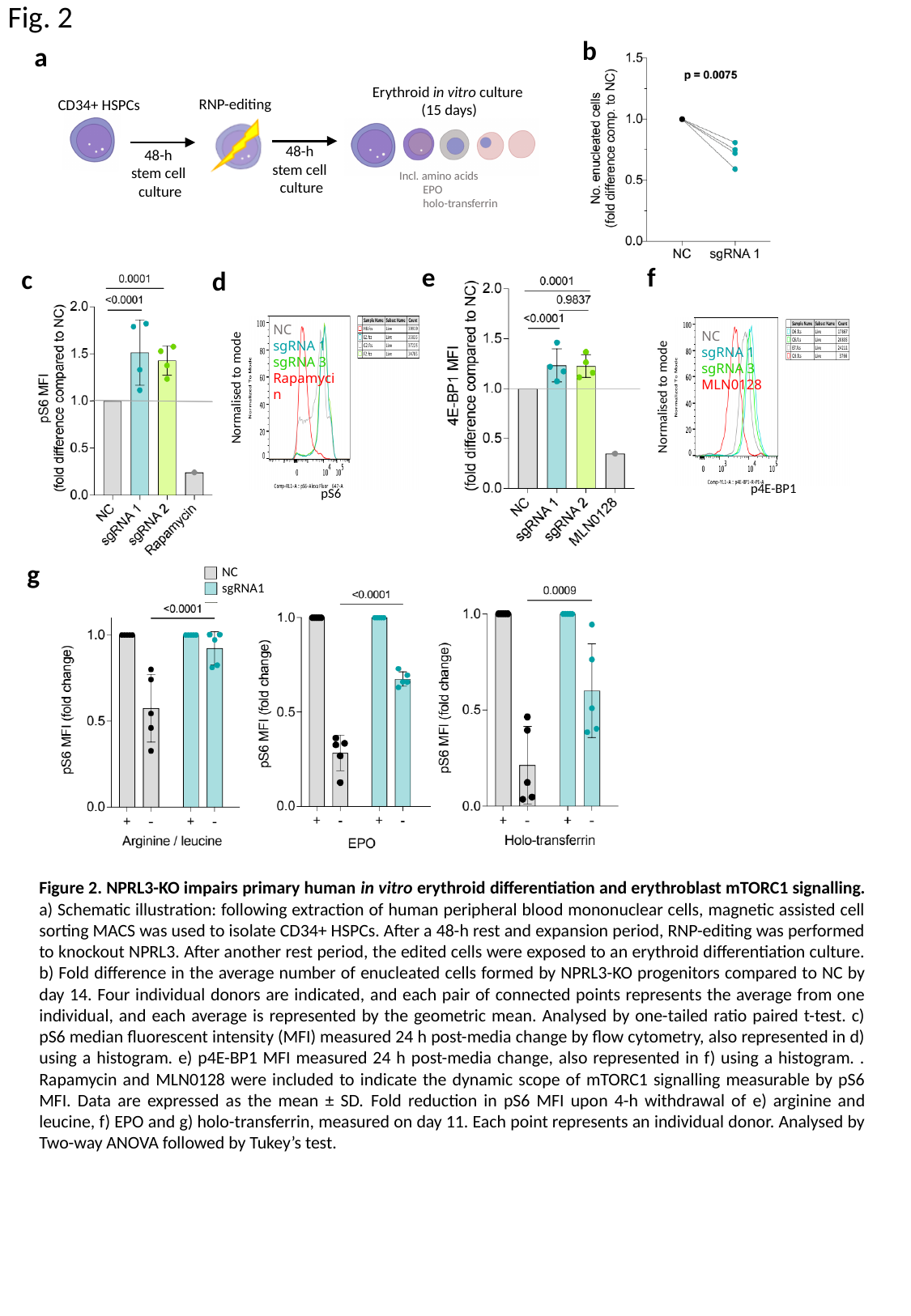

Fig. 2
b
a
Erythroid in vitro culture
(15 days)
RNP-editing
CD34+ HSPCs
48-h
stem cell
culture
48-h
stem cell
culture
Incl. amino acids
 EPO
 holo-transferrin
NC
sgRNA 1
sgRNA 3
Rapamycin
Normalised to mode
pS6
f
e
c
d
NC
sgRNA 1
sgRNA 3
MLN0128
Normalised to mode
p4E-BP1
g
NC
sgRNA1
Figure 2. NPRL3-KO impairs primary human in vitro erythroid differentiation and erythroblast mTORC1 signalling. a) Schematic illustration: following extraction of human peripheral blood mononuclear cells, magnetic assisted cell sorting MACS was used to isolate CD34+ HSPCs. After a 48-h rest and expansion period, RNP-editing was performed to knockout NPRL3. After another rest period, the edited cells were exposed to an erythroid differentiation culture. b) Fold difference in the average number of enucleated cells formed by NPRL3-KO progenitors compared to NC by day 14. Four individual donors are indicated, and each pair of connected points represents the average from one individual, and each average is represented by the geometric mean. Analysed by one-tailed ratio paired t-test. c) pS6 median fluorescent intensity (MFI) measured 24 h post-media change by flow cytometry, also represented in d) using a histogram. e) p4E-BP1 MFI measured 24 h post-media change, also represented in f) using a histogram. . Rapamycin and MLN0128 were included to indicate the dynamic scope of mTORC1 signalling measurable by pS6 MFI. Data are expressed as the mean ± SD. Fold reduction in pS6 MFI upon 4-h withdrawal of e) arginine and leucine, f) EPO and g) holo-transferrin, measured on day 11. Each point represents an individual donor. Analysed by Two-way ANOVA followed by Tukey’s test.

### Slide 4
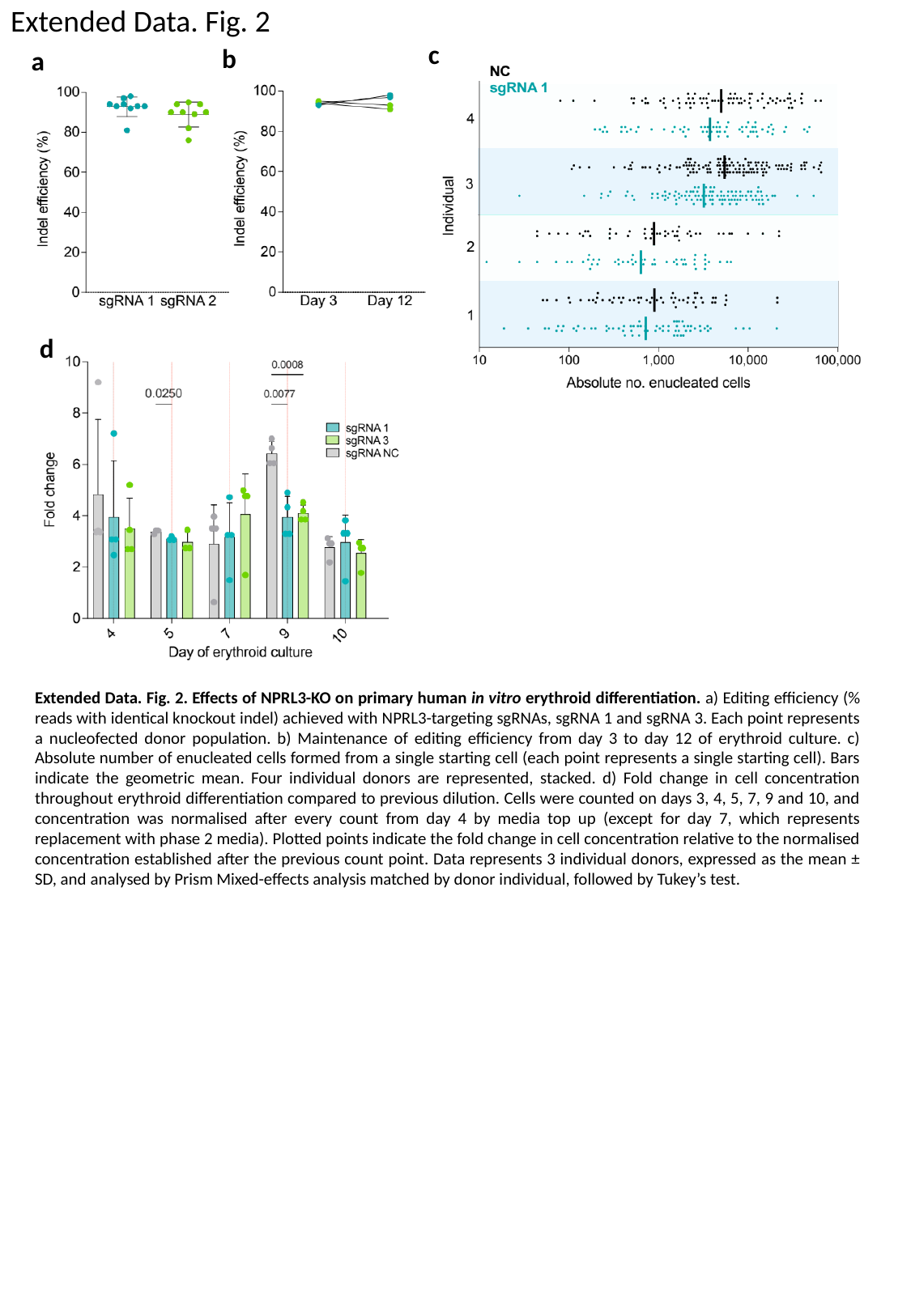

Extended Data. Fig. 2
c
b
a
d
Extended Data. Fig. 2. Effects of NPRL3-KO on primary human in vitro erythroid differentiation. a) Editing efficiency (% reads with identical knockout indel) achieved with NPRL3-targeting sgRNAs, sgRNA 1 and sgRNA 3. Each point represents a nucleofected donor population. b) Maintenance of editing efficiency from day 3 to day 12 of erythroid culture. c) Absolute number of enucleated cells formed from a single starting cell (each point represents a single starting cell). Bars indicate the geometric mean. Four individual donors are represented, stacked. d) Fold change in cell concentration throughout erythroid differentiation compared to previous dilution. Cells were counted on days 3, 4, 5, 7, 9 and 10, and concentration was normalised after every count from day 4 by media top up (except for day 7, which represents replacement with phase 2 media). Plotted points indicate the fold change in cell concentration relative to the normalised concentration established after the previous count point. Data represents 3 individual donors, expressed as the mean ± SD, and analysed by Prism Mixed-effects analysis matched by donor individual, followed by Tukey’s test.

### Slide 5
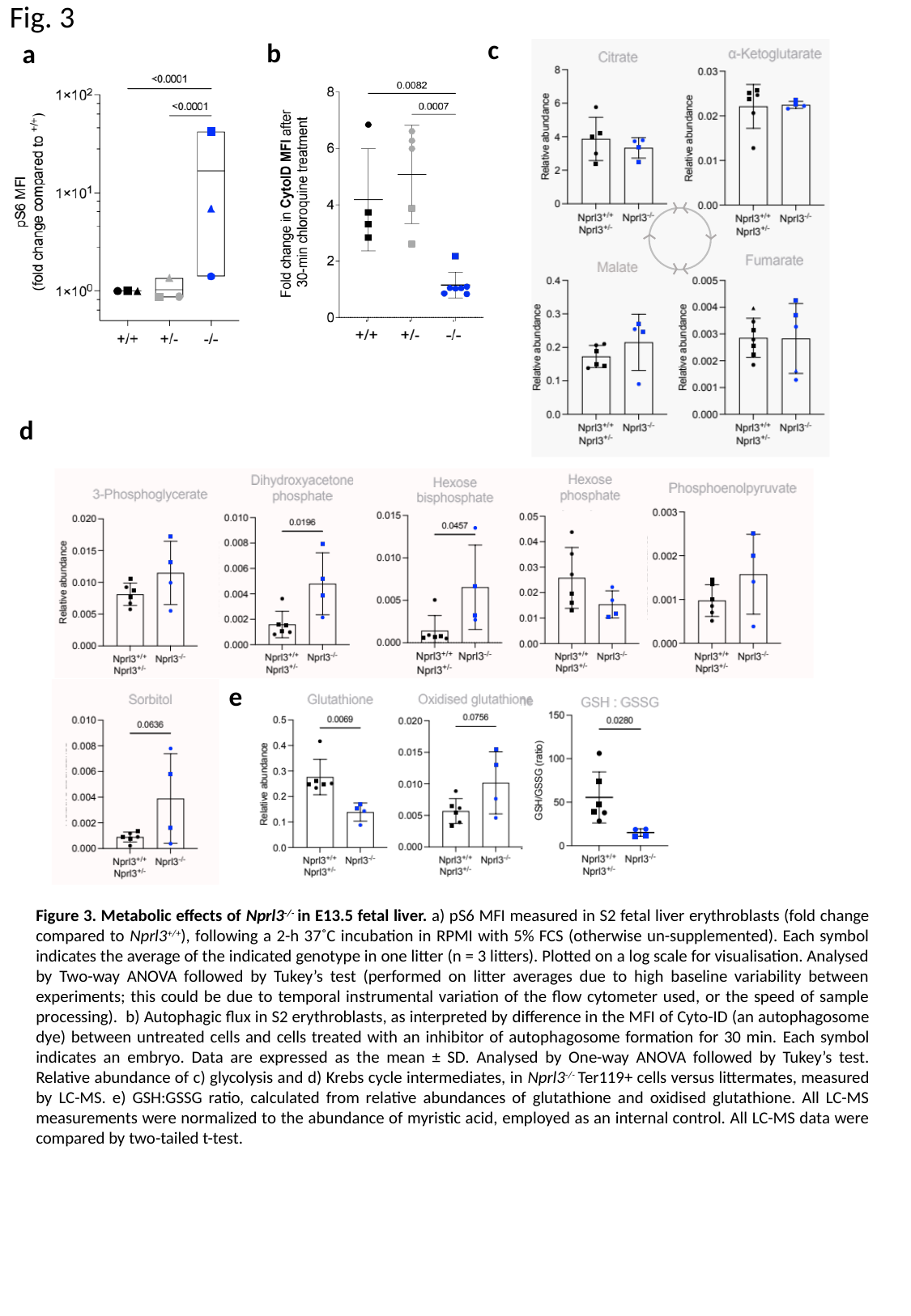

Fig. 3
c
b
a
d
e
Figure 3. Metabolic effects of Nprl3^-/-^ in E13.5 fetal liver. a) pS6 MFI measured in S2 fetal liver erythroblasts (fold change compared to Nprl3^+/+^), following a 2-h 37˚C incubation in RPMI with 5% FCS (otherwise un-supplemented). Each symbol indicates the average of the indicated genotype in one litter (n = 3 litters). Plotted on a log scale for visualisation. Analysed by Two-way ANOVA followed by Tukey’s test (performed on litter averages due to high baseline variability between experiments; this could be due to temporal instrumental variation of the flow cytometer used, or the speed of sample processing). b) Autophagic flux in S2 erythroblasts, as interpreted by difference in the MFI of Cyto-ID (an autophagosome dye) between untreated cells and cells treated with an inhibitor of autophagosome formation for 30 min. Each symbol indicates an embryo. Data are expressed as the mean ± SD. Analysed by One-way ANOVA followed by Tukey’s test. Relative abundance of c) glycolysis and d) Krebs cycle intermediates, in Nprl3^-/-^ Ter119+ cells versus littermates, measured by LC-MS. e) GSH:GSSG ratio, calculated from relative abundances of glutathione and oxidised glutathione. All LC-MS measurements were normalized to the abundance of myristic acid, employed as an internal control. All LC-MS data were compared by two-tailed t-test.

### Slide 6
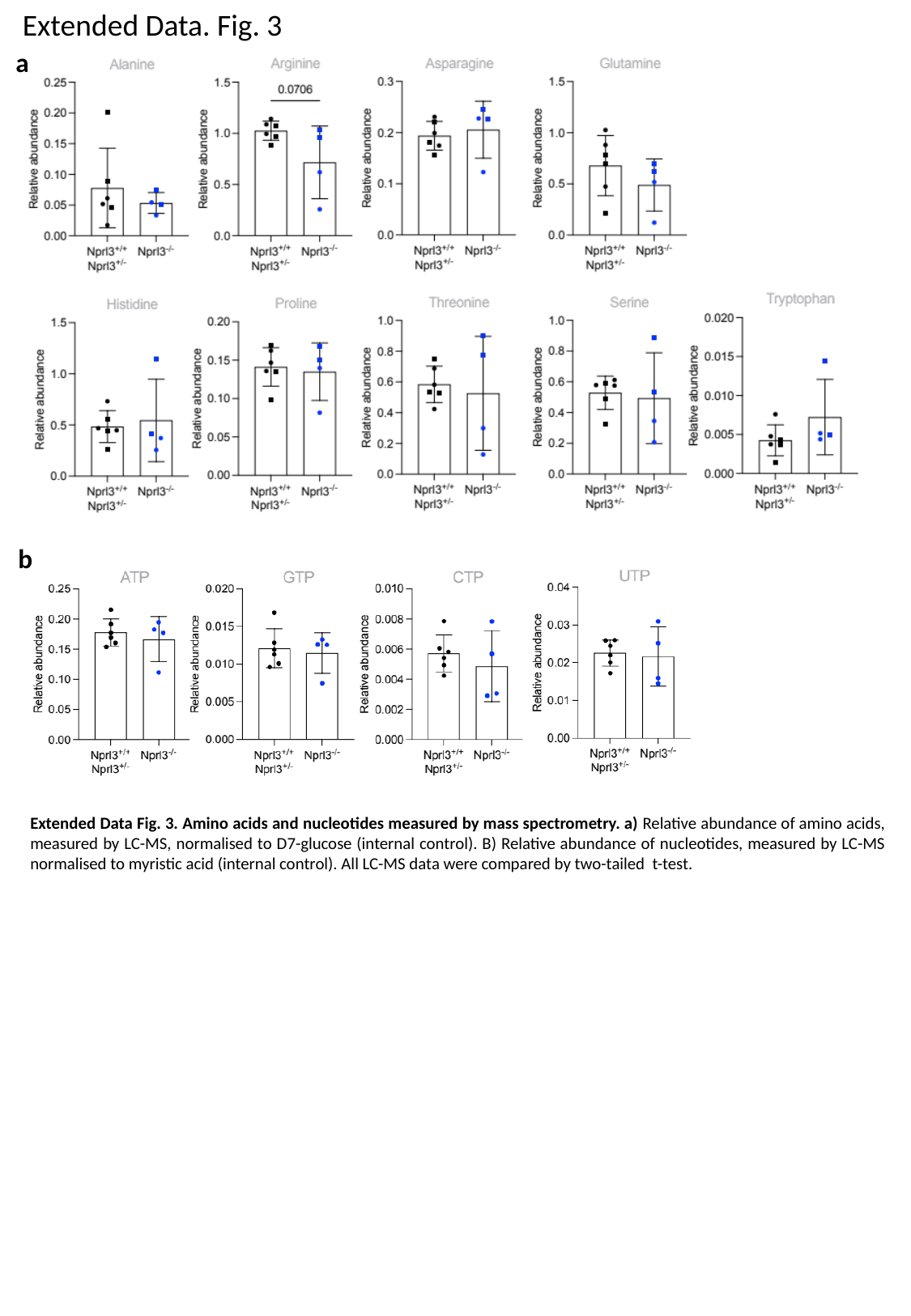

Extended Data. Fig. 3
a
b
Extended Data Fig. 3. Amino acids and nucleotides measured by mass spectrometry. a) Relative abundance of amino acids, measured by LC-MS, normalised to D7-glucose (internal control). B) Relative abundance of nucleotides, measured by LC-MS normalised to myristic acid (internal control). All LC-MS data were compared by two-tailed t-test.

### Slide 7
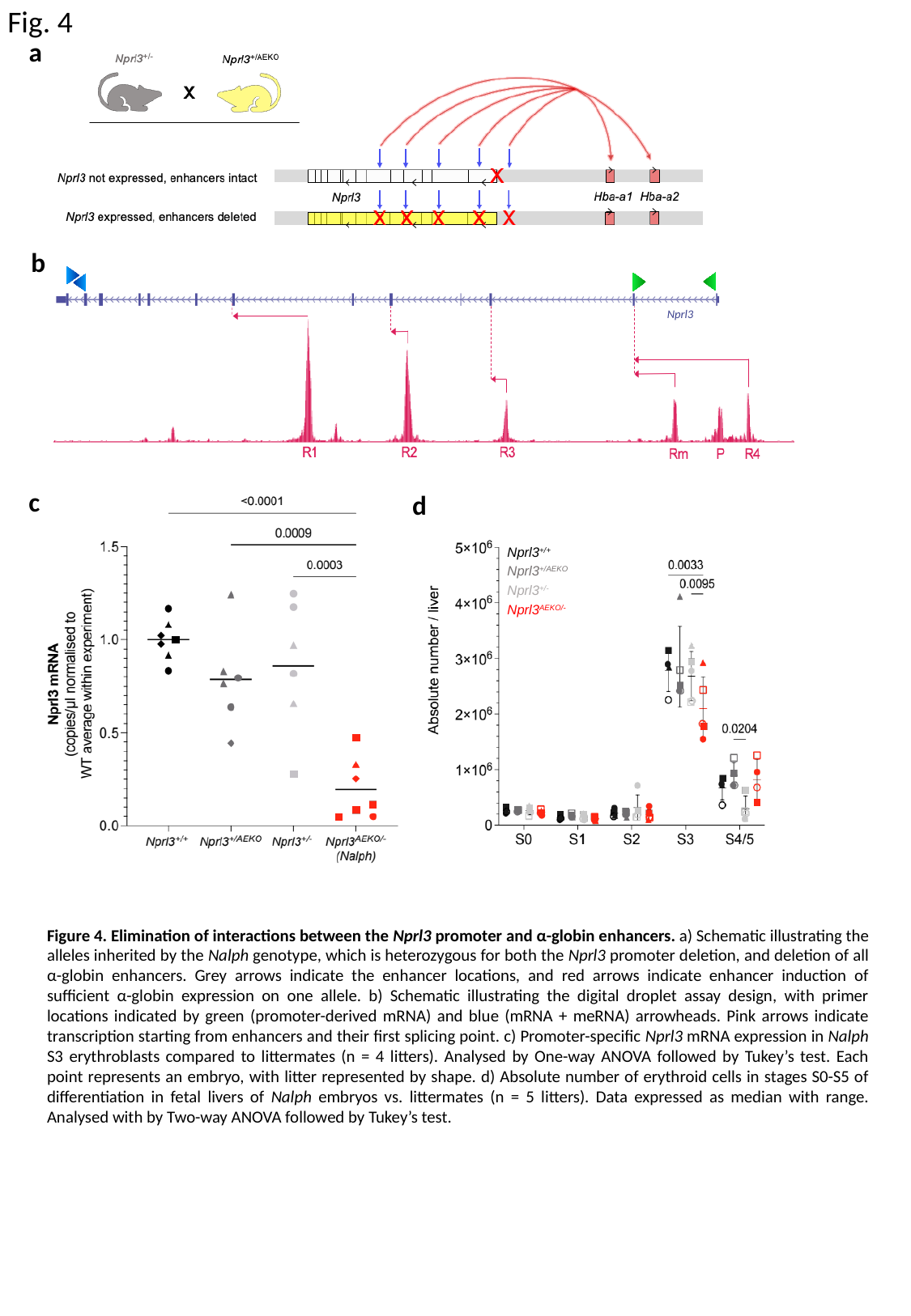

Fig. 4
a
b
Nprl3
c
d
Nprl3^+/+^
Nprl3^+/AEKO^
Nprl3^+/-^
Nprl3^AEKO/-^
Figure 4. Elimination of interactions between the Nprl3 promoter and α-globin enhancers. a) Schematic illustrating the alleles inherited by the Nalph genotype, which is heterozygous for both the Nprl3 promoter deletion, and deletion of all α-globin enhancers. Grey arrows indicate the enhancer locations, and red arrows indicate enhancer induction of sufficient α-globin expression on one allele. b) Schematic illustrating the digital droplet assay design, with primer locations indicated by green (promoter-derived mRNA) and blue (mRNA + meRNA) arrowheads. Pink arrows indicate transcription starting from enhancers and their first splicing point. c) Promoter-specific Nprl3 mRNA expression in Nalph S3 erythroblasts compared to littermates (n = 4 litters). Analysed by One-way ANOVA followed by Tukey’s test. Each point represents an embryo, with litter represented by shape. d) Absolute number of erythroid cells in stages S0-S5 of differentiation in fetal livers of Nalph embryos vs. littermates (n = 5 litters). Data expressed as median with range. Analysed with by Two-way ANOVA followed by Tukey’s test.

### Slide 8
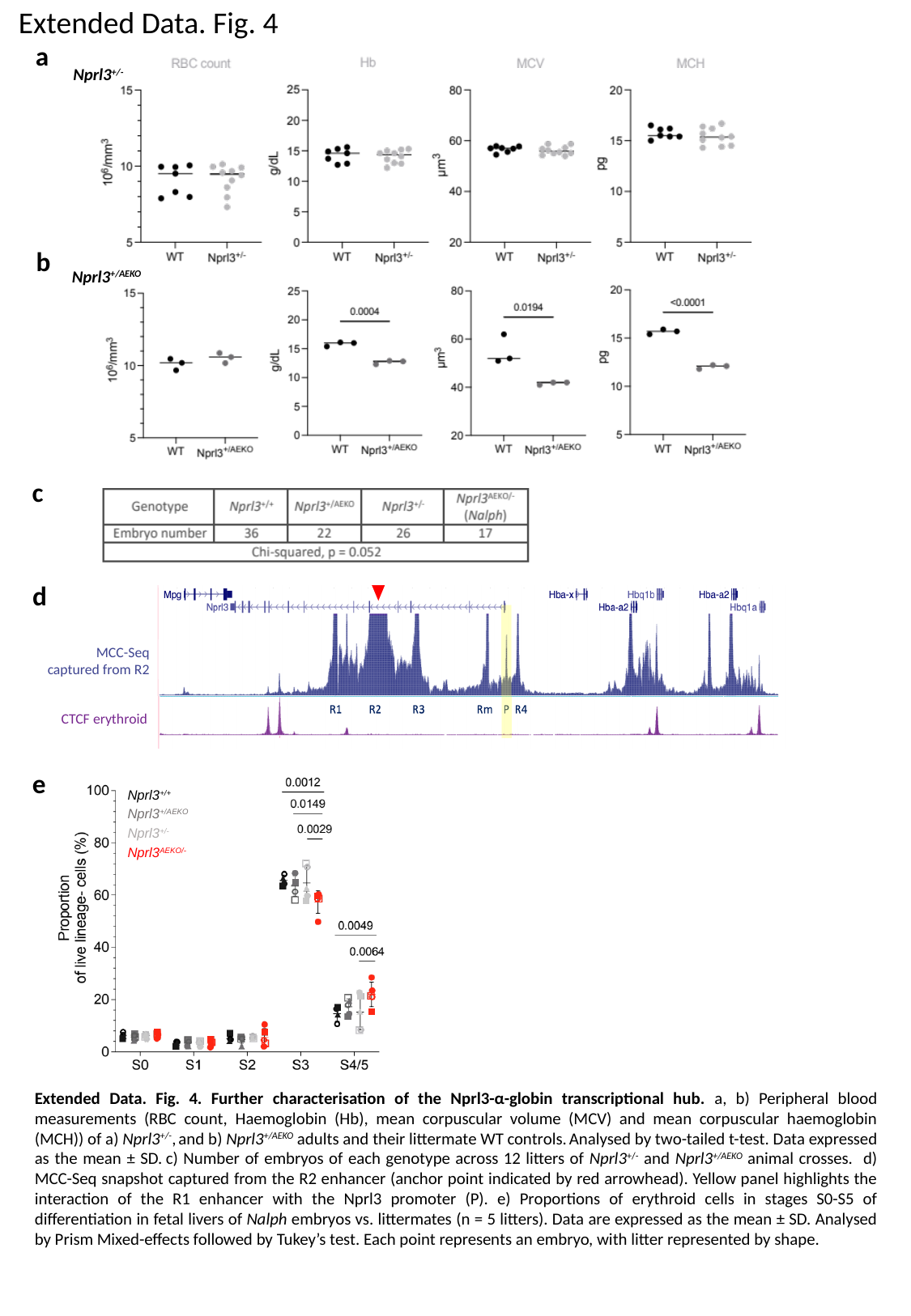

Extended Data. Fig. 4
a
Nprl3^+/-^
b
Nprl3^+/AEKO^
c
d
MCC-Seq
captured from R2
CTCF erythroid
e
Nprl3^+/+^
Nprl3^+/AEKO^
Nprl3^+/-^
Nprl3^AEKO/-^
Extended Data. Fig. 4. Further characterisation of the Nprl3-α-globin transcriptional hub. a, b) Peripheral blood measurements (RBC count, Haemoglobin (Hb), mean corpuscular volume (MCV) and mean corpuscular haemoglobin (MCH)) of a) Nprl3^+/-^, and b) Nprl3^+/AEKO^ adults and their littermate WT controls. Analysed by two-tailed t-test. Data expressed as the mean ± SD. c) Number of embryos of each genotype across 12 litters of Nprl3^+/-^ and Nprl3^+/AEKO^ animal crosses. d) MCC-Seq snapshot captured from the R2 enhancer (anchor point indicated by red arrowhead). Yellow panel highlights the interaction of the R1 enhancer with the Nprl3 promoter (P). e) Proportions of erythroid cells in stages S0-S5 of differentiation in fetal livers of Nalph embryos vs. littermates (n = 5 litters). Data are expressed as the mean ± SD. Analysed by Prism Mixed-effects followed by Tukey’s test. Each point represents an embryo, with litter represented by shape.

### Slide 9
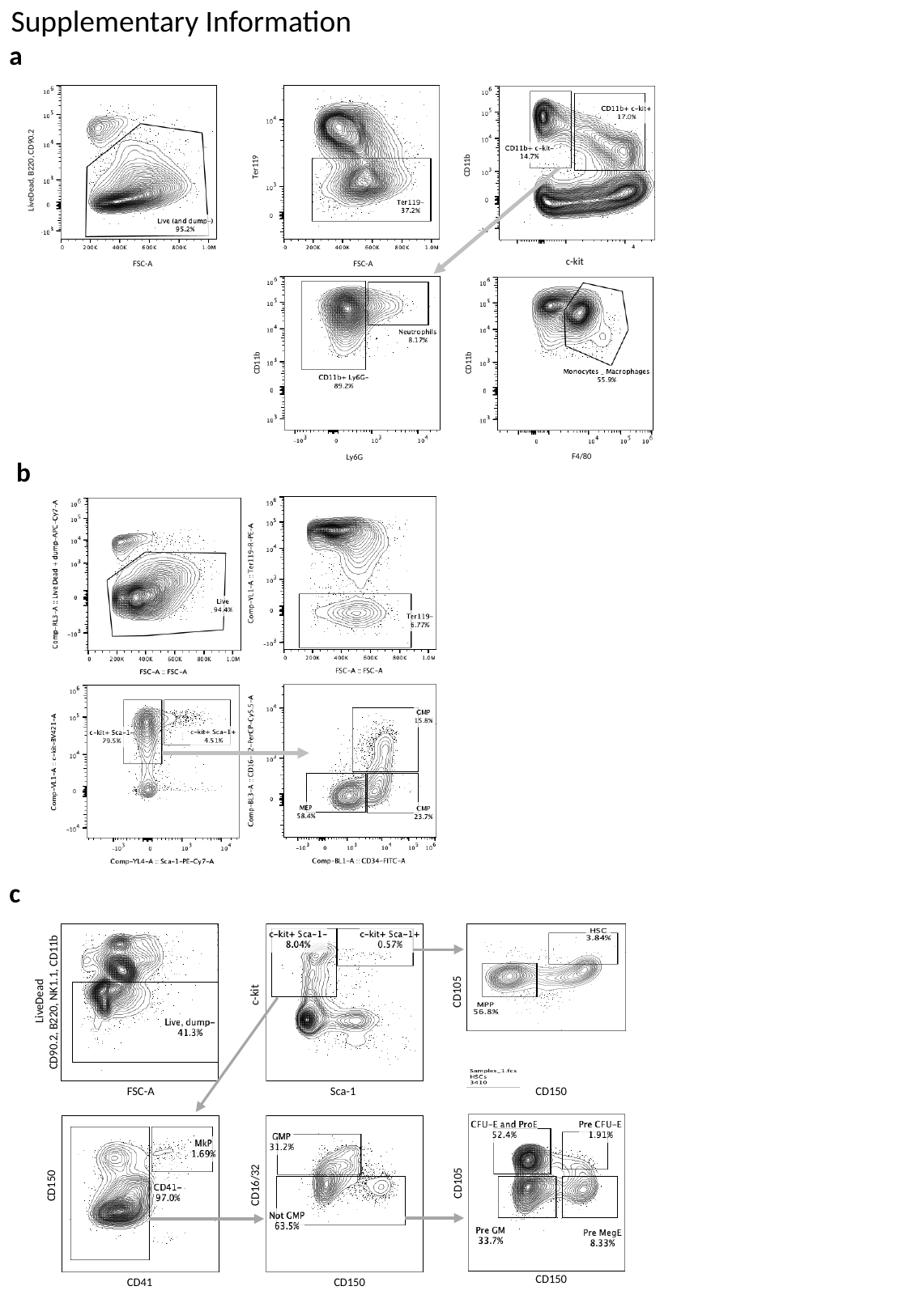

Supplementary Information
a
CD11b
Ter119
LiveDead, B220, CD90.2
c-kit
FSC-A
FSC-A
CD11b
CD11b
F4/80
Ly6G
b
c
c-kit
CD105
LiveDead
CD90.2, B220, NK1.1, CD11b
CD150
FSC-A
Sca-1
CD105
CD16/32
CD150
CD150
CD41
CD150
